# Cross-species analysis links cell-cell communication rewiring to NOTCH2 during serous endometrial carcinogenesis

**DOI:** 10.64898/2026.09.01.748540

**Authors:** Matalin G. Pirtz, Andrea Flesken-Nikitin, Coulter Q. Ralston, Daryl J. Phuong, Nating Wang, Tinyi Chu, Songru Chen, Yalin Zheng, Elisa Schmoeckel, Anna Yemelyanova, Ie-Ming Shih, Charles G. Danko, Benjamin D. Cosgrove, Alexander Yu. Nikitin

**Affiliations:** Department of Biomedical and Translational Sciences; Meinig School of Biomedical Engineering, Cornell University, Ithaca, New York, USA; Department of Genome Sciences, School of Medicine, University of Virginia, Charlottesville, Virginia, USA; Department of Pathology, Baltimore, MD, USA and Departments of Gynecology and Obstetrics, and Oncology, The Johns Hopkins School of Medicine, Baltimore, MD, USA; Institute of Pathology, School of Medicine and Health, Technical University of Munich, Munich, Germany; Department of Pathology & Laboratory Medicine, Weill Cornell Medicine, New York, New York, USA

**Author notes:** Correspondence to Alexander Yu. Nikitin.

## Abstract

Cell-cell interactions shape the fate of mutant cells during cancer initiation but how these interactions evolve during progression to pathologically recognizable lesions remain poorly understood. Here, we investigated cell-cell communication during serous endometrial carcinoma (SEC; also known as uterine serous carcinoma) development using a lineage-traceable mouse model and cross-species analyses of the mouse and human neoplastic endometrium. In mice, the early, pre-dysplastic stage was marked by a global decrease in inferred cell-cell interactions, followed by extensive communication network rewiring during neoplastic progression. Pathway-specific analysis revealed a similar pattern for NOTCH signaling, with NOTCH2 emerging as the dominant NOTCH receptor in *Trp53*/*Rb1*-mutant immature epithelial cells. Functionally, NOTCH2 promoted the outgrowth of more proliferative mutant organoids. Cross-species transcriptomic analysis identified conserved immature epithelial states in mouse and human neoplastic endometrial epithelium. In human tissues, NOTCH2 was overexpressed in serous endometrial intraepithelial carcinoma, a precursor of SEC, and in overt SEC. Furthermore, elevated NOTCH2 expression was associated with poor patient survival. These findings link cell-cell communication rewiring during experimental SEC development to conserved neoplastic epithelial states and identify NOTCH2 as an early marker and a potential target of disease interception.

## Introduction

Throughout life, normal tissues accumulate somatic mutations, as documented in the fallopian (uterine) tube ^1^, skin ^2^, esophagus ^3^, endometrium ^4^, and pancreas ^5^. Yet the prevalence of such alterations far exceeds the incidence of clinically apparent cancers, raising the question of which mechanisms restrain mutant cells and which enable their progression toward malignancy. Many mutant cells are likely constrained by interactions with their surrounding cellular niche, including cell-cell signaling ^6^. Understanding how mutant cells overcome these constraints and rewire local signaling networks during neoplastic progression is therefore critical for developing strategies for early cancer interception.

Such interception is particularly important for cancers in which early disease may remain clinically occult, including serous endometrial carcinoma (SEC; also known as uterine serous carcinoma). Uterine cancer is predicted to be the fifth leading cause of cancer-related death among women in the United States ^7^. In contrast to declining mortality from many other major cancers, both the incidence and mortality of uterine cancer have increased over the past four decades ^8^. Although SEC accounts for only approximately 10% of uterine cancers, it is responsible for about 40% of uterine cancer-related deaths ^9–11^. SEC is frequently diagnosed only after progression to advanced disease and is associated with high recurrence rates ^11,12^, contributing to its disproportionately high mortality.

SEC is characterized by the high frequency of *TP53* mutations and dysregulation of the RB-regulated cell-cycle pathway ^13–15^. These alterations are also characteristic of serous endometrial intraepithelial carcinoma (SEIC), a recognized precursor of SEC ^11,16–18^. However, how cells harboring these oncogenic alterations interact with their surrounding tissue environment during progression from histologically normal epithelium to recognizable neoplasia remains poorly understood.

Previously, we established a genetically engineered mouse model of SEC based on conditional inactivation of *Trp53* and *Rb1* in *Pax8*+ endometrial epithelial cells ^19^. Using single-cell and spatial transcriptomics together with immunohistochemistry, we subsequently identified diversification of the endometrial luminal epithelial compartment and acquisition of immature epithelial states during early, pre-dysplastic stages of SEC development ^20^.

Here, using this model, we define the dynamics of cell-cell communication during serous endometrial carcinogenesis and identify extensive signaling-network rewiring during neoplastic progression, with NOTCH signaling emerging as a prominent example of these changes. Cross-species transcriptomic analyses further identify conserved epithelial cell states in mouse and human endometrium. Finally, we establish the clinical relevance of NOTCH pathway alterations by showing that NOTCH2 is overexpressed in SEIC and SEC and that elevated NOTCH2 expression is associated with poor patient prognosis. Together, these findings link cell-cell communication rewiring during experimental SEC development to conserved epithelial states and clinically relevant NOTCH2 alterations in human serous endometrial carcinogenesis.

## Results

### Cell-cell interactions are rewired during early SEC development

We analyzed our previously published single-cell RNA-sequencing (scRNA-seq) dataset from a mouse model of serous endometrial carcinoma (SEC) at three stages of progression (Figure 1A). The integrated dataset included epithelial, fibroblast, endothelial, smooth muscle, and immune cell populations annotated as previously described ^20^ and (Supplemental Figure 1A-C).

**Figure 1.**
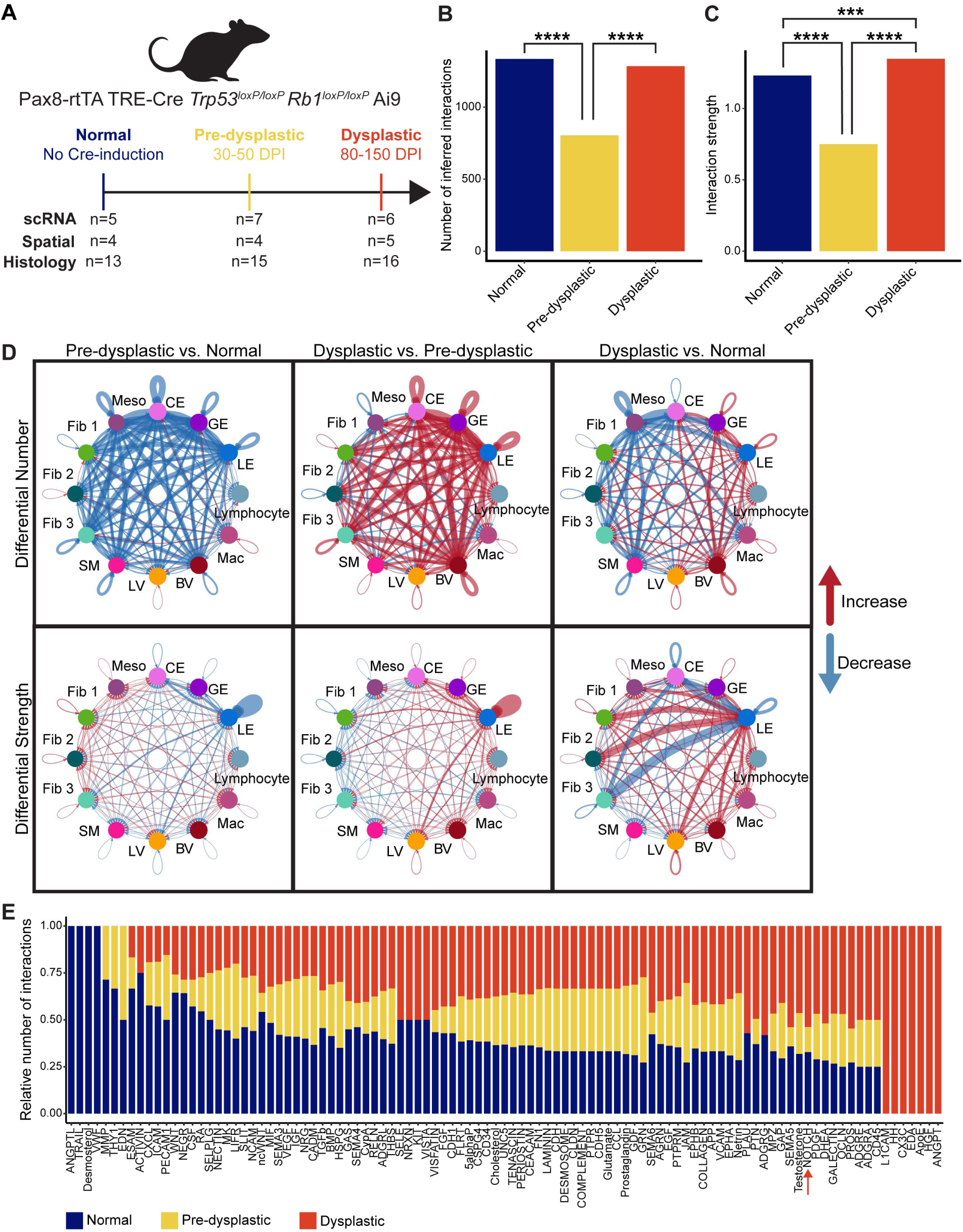
Cell-cell interaction changes during carcinogenesis. (A) Data collection timeline. Samples were collected at specific days post-induction (DPI) with a single intraperitoneal doxycycline injection for either single-cell RNA-sequencing, Visium RNA-sequencing, or paraffin-embedded for immunohistochemical analysis. n, number of samples per time point. (B and C) Predicted number (B) and strength (C) of interactions by stage. The strength of interactions is the summation of the probabilities of associated ligand-receptor pairs. ***P<0.001, ****P<0.0001, Fischer’s Exact tests. (D) Circle plots, differential interactions between clusters by stage. Thickness of connecting lines is proportional to the magnitude of differential number/strength of inferred interactions. Red connecting lines indicate an increase in interaction, whereas blue indicates a decrease relative to second listed stage at the top of columns. LE, Luminal Epithelium; GE, Glandular Epithelium; CE, Cycling Epithelium; Meso, Mesothelium; Fib, Fibroblast; SM, Smooth Muscle; LV, Lymphatic Endothelium; BV, Vascular Endothelium; Mac, Macrophage. (E) Relative contribution of each SEC stage to an individual CellChat annotated pathway based on number of predicted interactions. Arrow, highlighting the NOTCH pathway as one of interest.

To characterize changes in cell-cell communication during SEC development, we analyzed these data using CellChat ^21^, which infers intercellular signaling based on ligand-receptor expression. Pre-dysplastic samples exhibited a significant global reduction in both the predicted number and aggregate strength of cell-cell interactions (Figure 1B-C). In dysplastic samples, the number of inferred interactions recovered to near-normal levels, whereas overall interaction strength exceeded that observed in normal tissues, indicating substantial reorganization and enhancement of the predicted signaling network during progression.

Differential interaction analysis further showed that the pre-dysplastic stage was characterized by reduced communication among nearly all cell populations when compared to normal stage, with macrophages representing the major exception (Figure 1D). In contrast, both the number and strength of inferred interactions increased markedly in dysplastic relative to pre-dysplastic samples. This increase involved both epithelial-epithelial and epithelial-stromal communication, with fibroblast and endothelial interactions showing particularly prominent increases in dysplastic compared with normal tissues. Analysis of the relative contribution of each stage to individual signaling pathways revealed that many pathways followed this overall pattern (Figure 1E, Supplemental Figure 2A-B). Together, these findings identify a transient reduction in the native cell-cell communication network during the pre-dysplastic stage, followed by extensive signaling reorganization as histologically recognizable lesions emerge.

Several pathways, including NOTCH, CDH1, SEMA4, and LAMININ, followed this global trend, with the number and/or strength of inferred interactions decreasing during the pre-dysplastic stage and recovering during the dysplastic stage (Figure 1E, Supplemental Figure 2A). Among these, the NOTCH pathway was of particular interest because of its established roles in endometrial physiology and disease, including endometrial cancer ^22–25^. However, the specific contribution of NOTCH2 to SEC development remains poorly understood.

### NOTCH pathway interactions shift toward NOTCH2 during early SEC development

To define changes within the NOTCH signaling network, we examined the cell populations and ligand-receptor pairs contributing to inferred NOTCH interactions. To increase epithelial resolution, we transferred cell-state annotations from the stage-specific epithelial analysis reported previously ^20^. Luminal epithelial populations were then regrouped to generate a dataset containing luminal epithelial (LE), glandular epithelial (GE), DDP, COX/MAL, and cycling epithelial populations together with the stromal populations described above.

This analysis revealed marked simplification of the NOTCH interaction network during the pre-dysplastic stage compared with both normal and dysplastic tissues (Figure 2A), with a single fibroblast population, Fib2 becoming the predominant source of *Jag1*. By the dysplastic stage, the NOTCH signaling network regained complexity; however, the composition of signaling partners differed from that in normal tissues, with added fibroblast-mediated signaling and increased LE-derived signaling.

**Figure 2.**
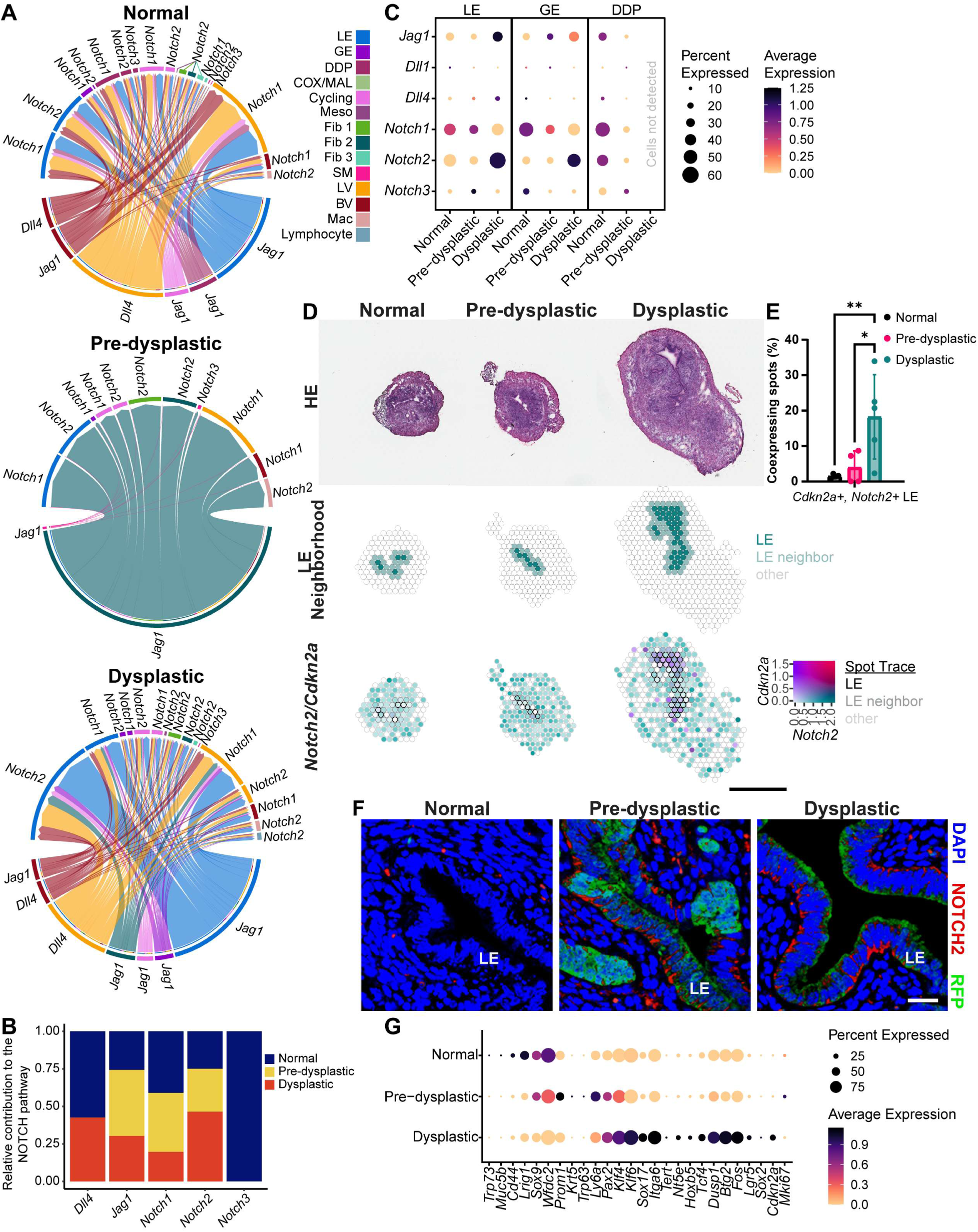
The NOTCH pathway undergoes remodeling in SEC. (A) Chord diagrams, predicted ligand-receptor pair interactions between cell types in normal (left), pre-dysplastic (center), and dysplastic (right) stages of SEC. Top arch, receptors; bottom arch, outer, ligands; bottom arch, inner, grouped receptors; colored by expressing cell type. Connecting line thickness indicates predicted strength/weight of the interaction. LE, Luminal Epithelium; GE, Glandular Epithelium; Meso, Mesothelium; Fib, Fibroblast; SM, Smooth Muscle; LV, Lymphatic Endothelium; BV, Vascular Endothelium. (B) Relative contribution of each NOTCH pathway component to the interaction network involving epithelial receiver cells to each stage based on weight. (C) Average log-normalized NOTCH pathway component expression in each epithelial population within each SEC stage. (D) Spatial feature plots, Hematoxylin and eosin (top), LE neighborhood (LE, LE neighbor, other), and colocalization of *Cdkn2a* with *Notch2*. Gene colocalization includes an LE neighborhood trace (black, LE; dark grey, LE neighbor; light grey, other). Scale bar, 2 mm for all images. (E) Quantification of *Notch2*, *Cdkn2a,* and luminal epithelial spots normalized to the number of spots coexpressing *Notch2* and *Cdkn2a* in spatial RNA sequencing samples. Normal (*n* = 4), pre-dysplastic (*n* = 4), and dysplastic (*n* = 5) spatial RNA-sequencing samples. *P < 0.05, **P < 0.01, two-way ANOVA, Tukey’s multiples test. Error bars denote SD. (F) Representative images of NOTCH2 (red) and RFP (green) expression in normal, pre-dysplastic, and dysplastic samples with DAPI (blue) counterstain. Confocal microscopy. Scale bar in bottom right image, 20 μm for all images. (G) Average log-normalized stem-like gene expression in the grouped LE, GE, and DDP populations in each SEC stage.

Notably, the relative contribution of epithelial interactions involving *Notch2* increased in both pre-dysplastic and dysplastic samples compared with normal tissues (Figure 2B and Supplemental Figure 2B). Thus, SEC progression was associated not only with changes in the overall extent of NOTCH communication but also with changes in the dominant receptor-ligand pairs contributing to the network. Pre-dysplastic samples showed prominent enrichment of *Notch1-Jag1* interactions. This pattern shifted in dysplastic samples, in which the relative contribution of *Notch1-Jag1* was lowest and interactions involving *Notch2*, particularly *Notch2-Jag1* and *Notch2-Dll4*, became more prominent (Supplemental Figure 2B).

### NOTCH2 expression increases during serous endometrial carcinogenesis

Consistent with the shift in inferred NOTCH interactions, epithelial *Notch2* expression was highest in dysplastic samples, with the strongest increase occurring within the LE population (Figure 2C), which we previously identified as a major contributor to SEC progression ^20^. In contrast, expression of canonical NOTCH target genes, including *Hes1* and *Hey1*, decreased during the pre-dysplastic stage and remained low in dysplastic tissues (Supplemental Figure 2C). The discordance between increased *Notch2* expression and reduced expression of these canonical downstream targets suggests that increased NOTCH2 expression during SEC development may not be accompanied by proportional activation of canonical NOTCH transcriptional output.

To determine the spatial relationship between *Notch2* expression and emerging mutant epithelial populations, we used a modified BayesPrism ^26^ framework, to deconvolve our Visium spatial RNA-sequencing data. LE regions were defined by histology and *Tacstd2* expression, allowing us to identify LE-enriched spots and evaluate their spatial relationship with other cell populations (Supplemental Figure 3A). As expected, predictions for LE and glandular epithelium (GE) were highest in histologically corresponding regions of normal and pre-dysplastic tissues. In dysplastic samples, LE-enriched spots became more spatially dispersed, consistent with the emergence of immature epithelial populations described previously ^20^.

We next quantified the spatial co-occurrence of *Notch2* with *Cdkn2a*, a marker associated with SEC in both mice and humans (Figure 2D-E) ^10,15,20^. *Cdkn2a/Notch2*-positive cells were significantly enriched within LE regions of dysplastic samples compared with normal tissues, consistent with the scRNA-seq findings.

NOTCH2 protein expression also increased during SEC development and was detectable at elevated levels beginning at the pre-dysplastic stage (Figure 2F and Supplemental Figure 3B-C). High NOTCH2 expression correlated with RFP lineage labeling (Figure 2F and Supplemental Figure 3B) and P16 expression (Supplemental Figure 3C), indicating preferential NOTCH2 upregulation in *Trp53/Rb1*-mutant LE cells. P16 frequently colocalized with high NOTCH2 expression; however, some P16-negative cells in pre-dysplastic tissues also showed elevated NOTCH2. Thus, increased NOTCH2 expression may identify a population of early mutated epithelial cells more inclusive than P16 alone.

As previously reported ^20^, dysplastic epithelial cells showed increased expression of genes associated with immature or stem-like states, including *Itga6, Klf4 and Sox2* (Figure 2G). These genes were preferentially expressed in cells with the highest levels of *Notch2* (Supplemental Figure 3D), linking elevated *Notch2* expression to the immature epithelial phenotype that emerges during SEC progression. *Cdkn2a* showed a similar association. In addition, *Mki67* expression was highest among *Notch2*-high cells during the pre-dysplastic stage, suggesting an association between elevated *Notch2* and early proliferative activity.

Together, these findings demonstrate increased NOTCH2 expression at both the transcript and protein levels in mutant LE cells during SEC development and link elevated NOTCH2 to immature and proliferative epithelial states.

### NOTCH2 overexpression selects for highly proliferative cells during SEC initiation

To determine whether increased NOTCH2 activity affects the growth of mutant endometrial epithelial cells, we established an endometrial epithelial organoid model of SEC (Figure 3A). Monoclonal organoids were generated from endometrial epithelial cells isolated from *Trp53^loxP/loxP^ Rb1^loxP/loxl^ Ai9* mice and transformed through Cre-mediated inactivation of *Trp53* and *Rb1* (Supplemental Figure 4A and ^27^).

**Figure 3.**
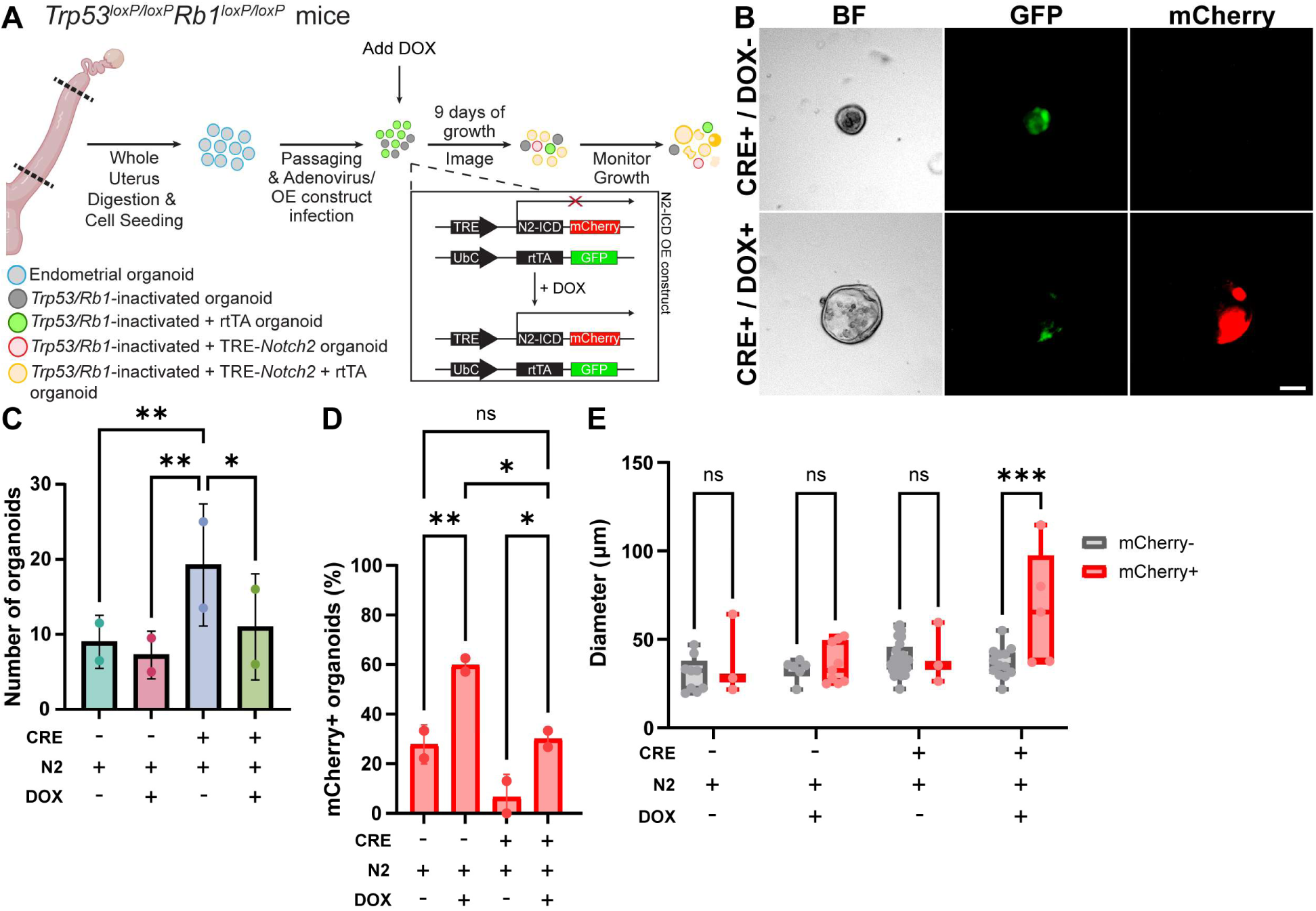
Overexpression of NOTCH2 affects the growth of mutant mouse endometrial organoids. (A) Schematic of the conditional overactivation of NOTCH2 in mutant mouse endometrial organoids. N2-ICD, NOTCH2 intracellular domain sequence. (B) Representative images of *Trp53* and *Rb1*-inactivated organoids with (bottom) or without (top) N2-ICD overexpression. Live fluorescence imaging. Scale bar in bottom right image, 20 μm for all images. (C) Organoid counts. (D) Percent of mCherry positive organoids relative to total organoids. (C-D) Each dot represents the average across two wells with two technical replicates per condition. (E) Comparison of mCherry-to mCherry+ organoid diameters within each condition. Each spot represents one organoid. (C-E) CRE, AdenoCre infection; N2, TRE-N2-ICD-mCherry + rtTA-GFP virus infection; DOX, doxycycline treated. Two-way ANOVA, Šídák’s multiple comparison, *P ≤ 0.05, **P ≤ 0.01, ***P ≤ 0.001. Error bars denote SD.

Conditional activation of NOTCH2 was achieved using a lentiviral reverse tetracycline transactivator (rtTA)/tetracycline response element (TRE) system to induce expression of the NOTCH2 intracellular domain (N2-ICD)-mCherry following doxycycline treatment (Figure 3A-B and Supplemental Figure 4B-C).

Induction of N2-ICD reduced the number of mutant organoids that formed (Figure 3C-D), indicating that high NOTCH2 activity decreased overall organoid-forming efficiency. However, the organoids that persisted following N2-ICD induction were significantly larger than those under all other experimental conditions (Figure 3E). These findings indicate that elevated NOTCH2 activity imposes a selective pressure on mutant epithelial cells while promoting robust expansion of the cells that tolerate or respond favorably to NOTCH2 activation. Thus, increased NOTCH2 activity during the pre-dysplastic stage may contribute to selection and expansion of highly proliferative mutant epithelial populations.

### Cross-species transcriptomic analysis supports a conserved role for NOTCH2 in SEC

To assess the relevance of our mouse findings to human SEC, we integrated publicly available human single-cell and single-nucleus RNA-sequencing datasets from Foley *et al.* ^28^, Garcia-Alonso *et al.* ^29^, Huang *et al.* ^30^, Lai *et al.* ^31^, Marečková *et al.* ^32^, and Wang *et al.* ^33^ (Supplemental Figure 5A-C). To maximize comparability with our diestrus-stage mouse model, epithelial and cycling cells from mice were initially compared with epithelial and cycling populations across human datasets.

Following preprocessing (Supplemental Figure 5B-C), the integrated human dataset contained 358,789 cells, including 176,357 epithelial cells identified by *EPCAM, KRT8,* and *PAX8* expression. Human epithelial cells were compared with mouse endometrial epithelial populations using Self-Assembling Manifold mapping (SAMap), an algorithm developed for cross-species transcriptomic comparisons ^34^. Human “Proliferative Early” and “Proliferative” phase epithelial populations showed the strongest correspondence with normal diestrus mouse epithelial cells (Supplemental Figure 5D). We therefore subsetted and reclustered cells from “Proliferative Early”, “Proliferative”, and “SEC” samples for subsequent analyses, yielding 163,003 cells (Supplemental Figure 6A). Clusters were annotated on the basis of previously defined expression profiles and differentially expressed genes (Methods and Supplemental Figure 6B-D). To minimize effects of patient heterogeneity and differences in sample collection (Supplemental Figure 6D-E), epithelial populations were consolidated into LE, GE, Hormone-Responsive Proliferative-like, Ciliated, and Cycling groups.

Mouse and human epithelial populations showed substantial correspondence in integrated transcriptomic space (Figure 4A-B). Human GE populations mapped most strongly to mouse GE, DDP, and COX/MAL populations, with some correspondence to mouse LE. Human LE, Hormone-responsive Progenitor-like, and Ciliated populations mapped exclusively to mouse LE (Figure 4C-D). These relationships were maintained across normal and SEC samples, supporting conservation of major epithelial cell states between the mouse model and human endometrium.

**Figure 4.**
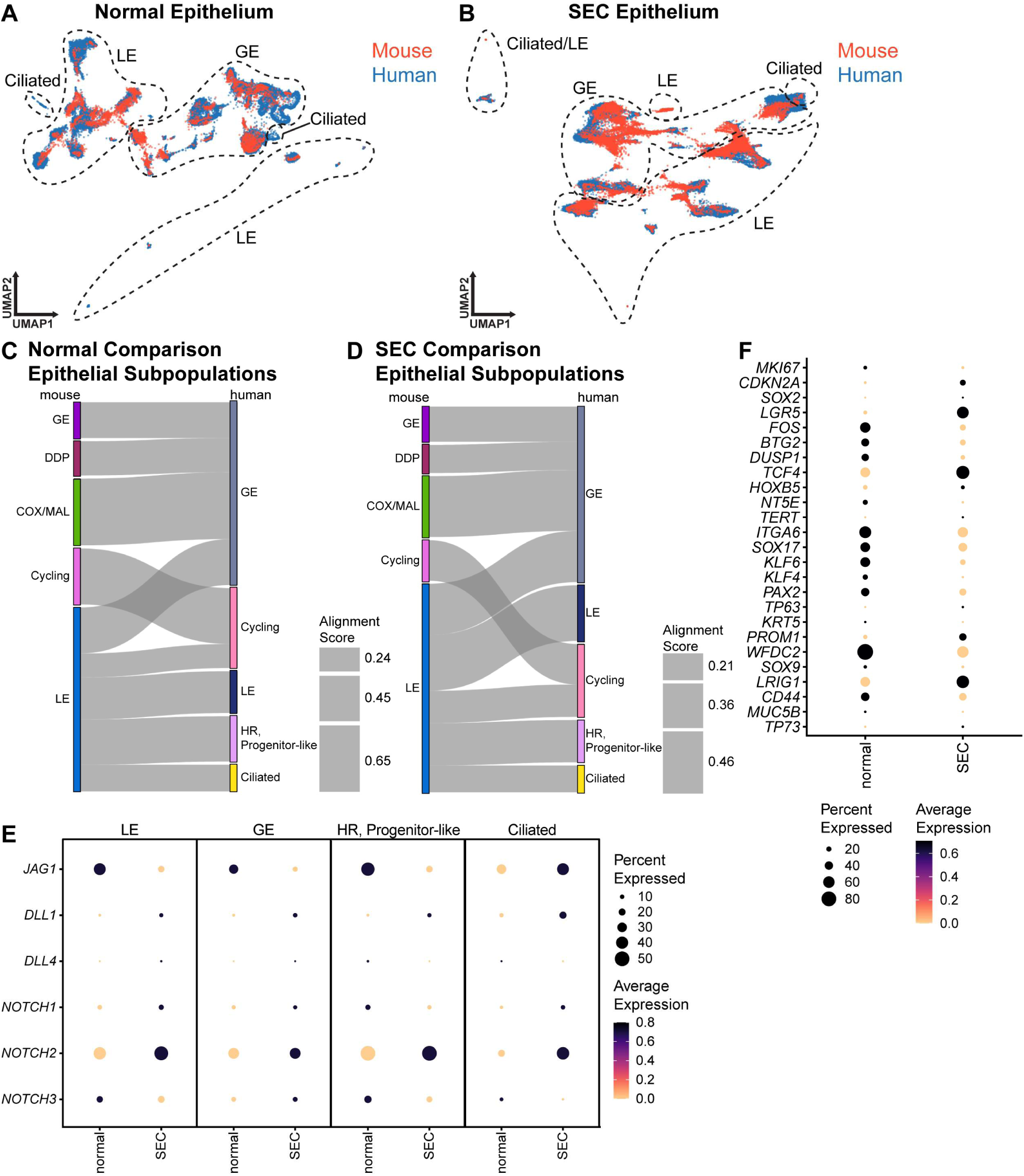
Cross-species analysis identifies conserved epithelial states and altered NOTCH and stem-like programs in SEC. (A-B) UMAP, SAMap comparison between mouse and human normal endometrial epithelium (A) and SEC endometrial epithelium (B), roughly grouped and labeled into general epithelial subtypes. (C-D) Sankey plots of the correlation between the human and mouse healthy (C) and SEC (D) endometrial epithelial subpopulations. Thickness of connecting lines proportional to the alignment score between human and mouse populations. (E) Average log-normalized NOTCH pathway component expression in LE, GE, Hormone-Responsive (HR) Progenitor-like, and Ciliated populations in human normal proliferative uterus samples and human SEC samples. (F) Average log-normalized stem-like gene expression in the grouped LE, GE, Hormone-Responsive Progenitor-like, and Ciliated populations in healthy and SEC samples.

We next compared expression of NOTCH pathway components between normal human endometrium and SEC. Among NOTCH receptors, *NOTCH2* showed consistently high expression across human SEC epithelial populations (Figure 4E), supporting conservation of the NOTCH2-associated phenotype observed in the mouse model. Consistent with the reduced canonical NOTCH transcriptional output observed in mice, canonical NOTCH target genes were preferentially expressed in normal rather than SEC epithelial populations (Supplemental Figure 7A).

Analysis of genes associated with immature or stem-like epithelial states further revealed increased *SOX2*, *LGR5,* and *CDKN2A* expression across human SEC epithelial populations, similar to the pattern observed in the mouse model. Expression of these genes was highest in *NOTCH2*-high cells (Figure 2G, Figure 4F, Supplemental Figures 3D and 7B).

### NOTCH2 is overexpressed in early human SEC lesions and is associated with poor patient survival

Consistent with the mouse model and human single-cell transcriptomic analyses, NOTCH2 expression, reflected by membranous and cytoplasmic staining, was significantly higher in SEC samples (n = 24) than in normal human endometrial epithelium and endometrioid endometrial carcinoma (EEC; n=21) (Figure 5A-B). We next assessed the clinical significance of *NOTCH2* expression using KMplotter ^35^. Among *CDKN2A*-high endometrial carcinoma cases, elevated *NOTCH2* mRNA expression was significantly associated with poorer survival (Figure 5C). Notably, *NOTCH2* was the only NOTCH receptor whose expression was significantly upregulated and associated with survival in this analysis (Supplemental Figure 8).

**Figure 5.**
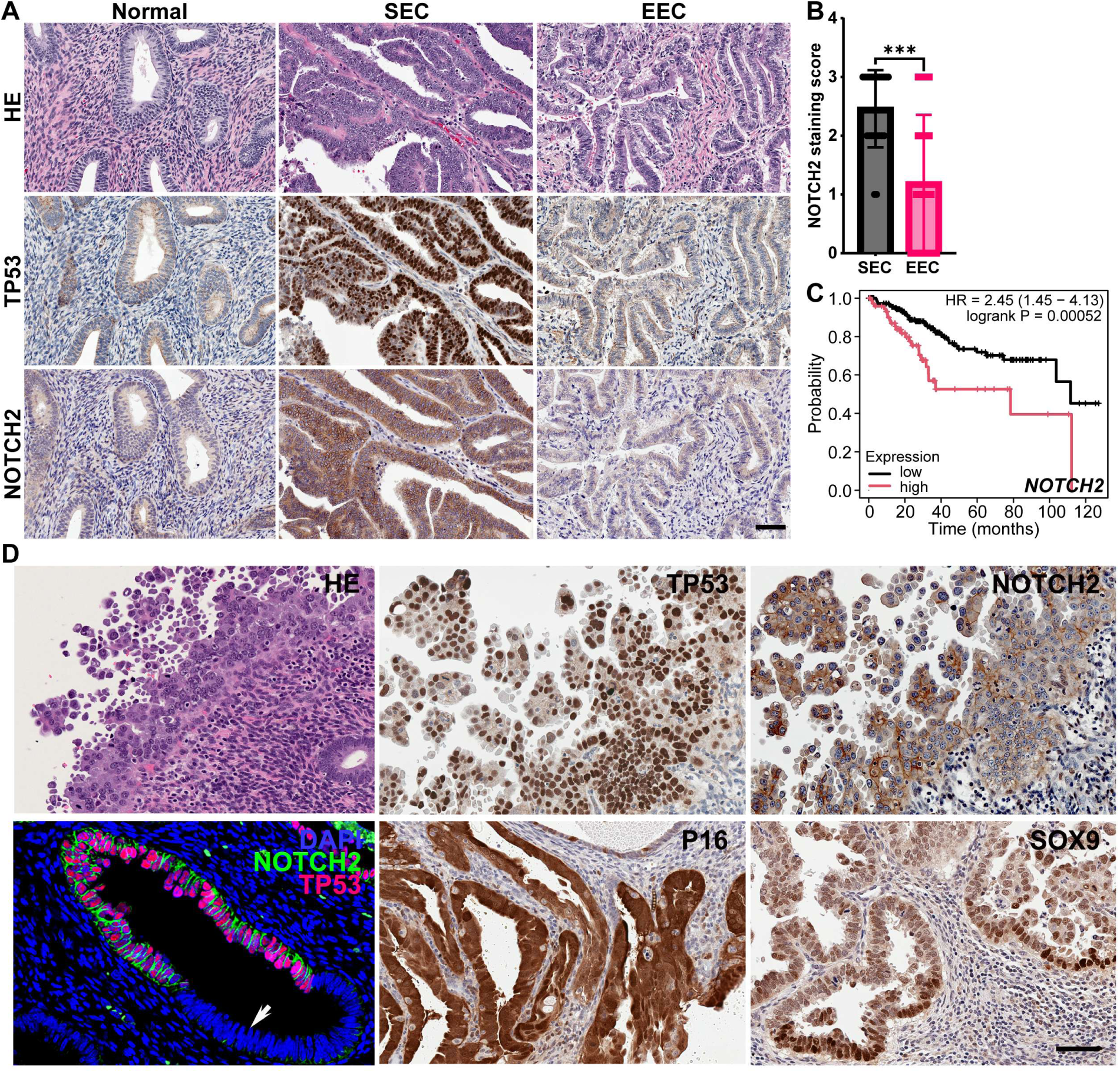
NOTCH2 overexpression characterizes human SEC and SEIC and is associated with poor patient survival. (A) Hematoxylin Eosin (HE) staining and immunohistochemical detection of TP53 and NOTCH2 (brown color) in endometrial epithelium (Normal), serous endometrial carcinoma (SEC) and endometrioid endometrial carcinoma (EEC). ABC Elite method with hematoxylin counterstaining. Scale bar, 60 μm for all images. (B) Quantification of NOTCH2 staining in SEC (*n*=24) and EEC (*n=*21). ***P≤ 0.001, Mann–Whitney test. Error bars denote SD. (C) Kaplan–Meier plots, Overall patient survival stratified by the mRNA expression of *NOTCH2*. Only patients with above median expression of CDKN2A were included (*n* = 271). (D) Immunohistochemical detection of TP53, NOTCH2, P16 and SOX9 in SEIC. Bottom left, double immunofluorescence (IF) demonstrating co-expression of NOTCH2 (green) and TP53 (red) in dysplastic epithelium but not in adjacent normal epithelium (arrow), with DAPI nuclear counterstaining (blue). All other images were obtained using the ABC Elite method with hematoxylin counterstaining. Scale bar, 60 μm for all images except the IF image (40 μm).

Importantly, high NOTCH2 expression was also detected in serous endometrial intraepithelial carcinoma (SEIC; n=17), a recognized precursor lesion of SEC, demonstrating that NOTCH2 upregulation is already present at an early, histologically identifiable stage of human serous carcinogenesis. Consistent with the mouse findings and previous reports, SEIC lesions also expressed P16 ^11,16–18^ and the progenitor-associated marker SOX9 (Figure 5D). Together, these findings suggest that NOTCH2 may have utility as an auxiliary marker of early serous lesions and as a potential target for early SEC interception.

Taken together, these findings support the clinical relevance of NOTCH2 upregulation during serous endometrial carcinogenesis.

## Discussion

Here, we define changes in cell-cell communication accompanying the earliest stages of serous endometrial carcinoma (SEC) development and identify NOTCH2 as a prominent component of this process. Using a temporally controlled *Trp53/Rb1*-deficient mouse model, we found that the pre-dysplastic stage is characterized by a broad decrease in inferred cell-cell interactions, followed by extensive remodeling of the communication network as dysplasia develops. Within this changing landscape, NOTCH signaling underwent a particularly striking reorganization, with NOTCH2 emerging as the dominant NOTCH receptor in mutant immature epithelial cells. Functional studies further supported a role for NOTCH2 in promoting the outgrowth of proliferative mutant epithelial populations. Cross-species analyses identified related immature epithelial states in mouse and human neoplastic endometrium, while human tissue analyses demonstrated increased NOTCH2 expression in SEIC and overt SEC and an association between elevated NOTCH2 and adverse clinical outcome. Together, these findings suggest that early SEC development involves not simply progressive activation of intercellular signaling, but a transition from disrupted normal communication networks to the establishment of tumor-associated interactions favoring specific mutant epithelial cell states.

One of the most prominent features of early SEC development was the global reduction in inferred cell-cell interactions during the pre-dysplastic stage. This change coincided with the emergence of distinct luminal epithelial (LE) populations previously identified in this model ^20^. Rather than showing a gradual increase in tumor-promoting communication following *Trp53/Rb1* loss, the developing neoplastic tissue therefore passes through a period of reduced connectivity before establishing a remodeled network in dysplastic lesions, including increased interactions involving epithelial, fibroblast, and endothelial populations. These observations raise the possibility that disruption of homeostatic communication represents an early feature of mutant epithelial cell adaptation and creates a permissive context for subsequent network remodeling. Determining whether this transient reduction in communication is required for progression, or instead reflects the changing composition and state of the tissue, will require direct functional interrogation of the implicated interactions.

Among the pathways exhibiting this pattern, NOTCH signaling was of particular interest because of its established roles in endometrial regeneration ^36,37^ and endometrial carcinoma ^22–24^, yet its specific contribution to SEC remains poorly defined. This distinction is important because NOTCH signaling can exert either tumor-promoting or tumor-suppressive effects depending on cellular context and on the receptors and ligands involved ^38,39^. In our model, the pre-dysplastic stage was characterized by marked simplification of the NOTCH interaction network, including loss of several epithelial interactions present in normal endometrium. This was followed by establishment of a distinct, NOTCH2-dominant network during dysplastic progression. Importantly, NOTCH2-associated epithelial interactions began to increase during the pre-dysplastic stage, and *Notch2* expression progressively increased in mutant LE cells at both the spatial transcriptomic and protein levels. Thus, NOTCH2 activation occurs within the period in which the normal NOTCH network is being dismantled and precedes establishment of the dysplastic interaction network.

Our functional findings further suggest that increased NOTCH2 signaling alters the fitness of emerging mutant epithelial populations. Cells with high *Notch2* expression exhibited increased *Mki67* expression, and increased NOTCH2 activation favored the outgrowth of more proliferative mutant organoids. These findings are consistent with a cell-state–dependent function of NOTCH2 during SEC initiation rather than a uniform proliferative effect across the mutant epithelium. One possibility is that NOTCH2 activation promotes differentiation or restricts proliferative potential in some epithelial states while favoring the selection or expansion of mutant populations that remain highly proliferative. Such context dependence would be consistent with the diverse and sometimes opposing functions attributed to NOTCH signaling in cancer. More broadly, our observations support a model in which the consequences of NOTCH signaling during SEC progression depend on both the stage of transformation and the specific ligand-receptor and cellular context.

Cross-species transcriptomic comparisons further connected the NOTCH2-associated phenotype in the mouse model with epithelial states present in human SEC. In mice, progression was accompanied by expansion of immature epithelial phenotypes, and *Notch2*-high dysplastic cells showed increased expression of stem/progenitor-associated genes, including *Sox2, Itga6,* and *Lgr5*. Although these associations do not establish that NOTCH2 directly induces a stem-like state, they link increased *Notch2* expression with the immature epithelial program that becomes prominent during SEC progression. Notably, *SOX2* and *LGR5* showed similar patterns in human SEC. This finding is consistent with previous observations that LGR5 is associated with increased proliferation and adverse outcome in endometrial carcinoma ^40^, whereas SOX2-expressing endometrial cancer cells display greater colony-forming capacity than their SOX2-negative counterparts ^41^. Thus, the cross-species analyses suggest that the epithelial states associated with NOTCH2 activation in the mouse model capture features of immature epithelial populations present in human disease.

The human findings provide additional clinical relevance to this model. Increased NOTCH2 expression was observed in SEC and was associated with poor clinical outcome. Importantly, NOTCH2 overexpression was already evident in serous endometrial intraepithelial carcinoma (SEIC), a recognized precursor of SEC, indicating that increased NOTCH2 is acquired early rather than being restricted to advanced tumors. These observations raise the possibility that NOTCH2 could serve as an adjunctive marker for early serous lesions and assist in distinguishing SEC from high-grade endometrioid endometrial carcinoma when interpreted together with histopathologic and established molecular features. Validation in independent cohorts, including assessment of sensitivity and specificity in diagnostically challenging lesions, will be required before such an application can be established.

The findings of the current study further underscore the importance of NOTCH signaling in the pathogenesis of gynecologic serous carcinomas, while suggesting that distinct NOTCH receptor programs may operate according to the site of origin. In contrast to the prominent role of NOTCH2 identified here in SEC, NOTCH3 has been implicated in tubo-ovarian high-grade serous carcinoma (HGSC). Amplification within *NOTCH3* locus has been reported in HGSC, and activation of the NOTCH3-JAG-PBX1 axis promotes tumor growth ^42–44^. NOTCH3 overexpression has also been associated with ovarian cancer recurrence and resistance to carboplatin ^45^. Together with our findings, these observations suggest that uterine and tubo-ovarian serous carcinomas, despite shared histopathologic and molecular features, may preferentially engage distinct NOTCH receptor–dependent programs during tumor development and progression.

The preferential involvement of distinct NOTCH receptors in different serous carcinoma contexts may also have important therapeutic implications. Multiple strategies have been developed to therapeutically target NOTCH signaling in cancer, including inhibition at the level of ligand-receptor interactions, receptor cleavage, and downstream transcriptional activation ^46^. Tarextumab (OMP-59R5), a monoclonal antibody targeting NOTCH2 and NOTCH3, demonstrated antitumor activity in preclinical models of multiple epithelial malignancies, including ovarian cancer ^47^, and subsequently entered clinical testing. Although a randomized phase II trial in pancreatic cancer failed to demonstrate clinical benefit ^48^, these studies established the feasibility of receptor-directed NOTCH2/3 inhibition and emphasized the importance of tumor context and biomarker-based patient selection. More broadly, the clinical activity and FDA approval of the γ-secretase inhibitor nirogacestat in progressing desmoid tumors demonstrates that pharmacologic inhibition of NOTCH pathway activation can produce meaningful clinical benefit, although this agent is neither NOTCH2-selective nor directly informative regarding SEC. Our findings identify p53/RB1-deficient SEC as a biologically defined setting in which receptor-selective NOTCH2 inhibition warrants further investigation. In particular, determining whether NOTCH2-high SEC cells are selectively dependent on NOTCH2 signaling will be important for evaluating the therapeutic potential of this approach.

The strong association of NOTCH2 with p53-abnormal disease also suggests a potential mechanistic connection between p53 loss and remodeling of NOTCH signaling. p53 can directly regulate *NOTCH1* transcription ^49^, whereas regulation of NOTCH2 may occur indirectly through p53-responsive microRNAs. Among these, *miR-34b* is transcriptionally activated by p53 ^50^ and has been reported to directly target *NOTCH2* in ovarian cancer cells, where restoration of *miR-34b* reduced NOTCH2 expression, proliferation, and epithelial-to-mesenchymal transition ^51^. Loss of functional p53 could therefore contribute to increased NOTCH2 expression through loss of miR-34–mediated repression and potentially favor a shift from the normal NOTCH receptor network toward NOTCH2-dominant signaling. This mechanism remains speculative in SEC and will require direct experimental testing, but it provides a plausible link between one of the defining molecular alterations of SEC and the NOTCH2 phenotype identified here.

Interestingly, increased NOTCH2 expression was not accompanied by a corresponding increase in epithelial canonical NOTCH target gene expression in either species. This apparent discordance indicates that increased receptor abundance cannot be equated with uniform activation of the canonical NOTCH transcriptional program. Several possibilities could account for this observation, including transient or cell-state– restricted canonical signaling or engagement of noncanonical NOTCH functions. Noncanonical NOTCH signaling can occur through ligand-independent, cleavage-independent, or CSL-independent mechanisms, and interactions between NOTCH proteins and NF-κB, mTORC, PTEN/AKT, Wnt, Hippo, and TGF-β pathways have been implicated in cancer ^52,53^. Our findings therefore raise, but do not establish, the possibility that noncanonical NOTCH2 signaling contributes to SEC progression. Defining the relative contributions of canonical and noncanonical NOTCH2 activity will be an important direction for future studies and may be particularly relevant to the development of receptor- or pathway-selective therapeutic strategies.

Several limitations should be considered when interpreting the cross-species analyses. The human single-cell data were constrained by the publicly available datasets, and an appropriate healthy postmenopausal endometrial sc/snRNA-seq reference was not available. Because NOTCH signaling contributes to epithelial cell-state transitions in the cycling endometrium ^29^, comparison of SEC with cycling normal endometrium may obscure disease-associated differences in individual NOTCH pathway components and other markers, such as *SOX9*. Comparisons of stem/progenitor-associated programs may be similarly affected. In addition, computationally inferred ligand-receptor interactions identify candidate communication relationships rather than demonstrating physical or functional signaling between cell populations. These limitations emphasize the importance of the cross-species, spatial, protein, and functional approaches used here, while also identifying areas in which analysis of well-annotated human precursor lesions and postmenopausal normal endometrium will be particularly informative.

Although our study focused on NOTCH2, the broader communication analysis identified additional pathways that may participate in SEC initiation. SEMA4 signaling has been associated with ovarian carcinoma progression and proliferation ^54,55^, whereas altered LAMININ signaling has been linked to endometrial carcinoma severity and cell motility ^56^. CDH1 is required for normal endometrial regeneration and glandular organization ^57^, and combined *Trp53* and *Cdh1* inactivation produces aggressive endometrial tumors accompanied by alterations in the tumor microenvironment ^58^. We also identified pathways emerging preferentially during dysplasia, including ANGPT signaling, which has been implicated in endometrial carcinoma angiogenesis, immune infiltration, and outcome ^59,60^. These pathways provide additional candidates for understanding how early disruption of homeostatic signaling is converted into the multicellular communication network characteristic of overt neoplasia.

Collectively, our findings identify remodeling of cell-cell communication as an early feature of SEC development and place NOTCH2 at the intersection of mutant epithelial cell-state transitions, proliferative fitness, and progression to histologically recognizable disease. The detection of increased NOTCH2 in SEIC, its association with p53-abnormal endometrial carcinoma and adverse outcomes, and the conservation of related immature epithelial states between mouse and human support the relevance of this pathway to human serous endometrial carcinogenesis. More broadly, these findings suggest that the transition from mutant but morphologically normal epithelium to overt neoplasia may depend not only on cell-intrinsic genetic alterations, but also on the selective dismantling and reconstruction of the communication networks that determine mutant cell fate.

## Methods

### Human tissue acquisition

De-identified sectioned slides of the endometrium of patients with or without SEC, SEIC, or EEC were collected from women aged 42 - 86 years and prepared for histology at Weill Cornell Medicine and the Johns Hopkins School of Medicine. Tissue microarray of endometrial carcinoma and normal endometrial tissue, with grade data, was obtained from Quickarrays Inc (Fairfield, CA, USA; Cat No. EMC1021).

### Experimental animals

B6.Cg-Tg(Pax8-rtTA2S*M2)1Koes/J (Pax8-rtTA mice; RRID:IMSR_JAX:007176), B6.Cg-Tg(tetO-Cre)1Jaw/J (Tre-Cre mice; RRID:IMSR_JAX:006234) and *B6.Cg-Gt(ROSA)26Sor^tm9(CAG–tdTomato)Hze^* (Ai9 mice; RRID:IMSR_JAX:007909) were obtained from The Jackson Laboratory (Bar Harbor, ME, USA). *Trp53^loxP/loxP^* and *Rb1^loxP/loxP^* mice, which have *loxP* alleles flanking their respective genes were a gift from Dr. Anton Berns (The Netherlands Cancer Institute, Amsterdam, The Netherlands). For all experiments, mice were collected in late diestrus/early proestrus as previously described ^61–64^ and in Supplemental Methods.

### Doxycycline induction

Doxycycline was administered via a single intraperitoneal injection to Pax8-rtTA Tre-Cre *Trp53^loxP/loxP^Rb1^loxP/loxP^* Ai9 mice and control mice at 6–24 weeks old, as previously described^19,20^.

### Pathological evaluation

All mice underwent gross pathology evaluation at the time of necropsy, followed by histology, immunohistochemistry, and image analysis as previously described^20^. Dysplastic samples were diagnosed based on atypical nuclear to cytoplasmic ratios, mitotic figures, and atypical glandular patterning in the endometrial epithelium, as previously described^20^. Pre-dysplastic samples lacked morphologically discernable neoplastic lesions but were marked by presence of tdTomato/RFP staining indicative of Cre-*LoxP* mediated recombination. All human cases underwent standardized pathology review and H-score semi-quantification. All primary antibodies used for immunostaining are listed in Supplemental Table 1.

### Single-cell RNA-sequencing library preparation

All mouse single-cell RNA-sequencing (scRNA-seq) samples used in this manuscript were from previously reported transcriptomes^20^. As such, library preparation has been previously reported and data deposited in the Gene Expression Omnibus (GEO) under the accession code GSE269332.

### Download and alignment of human RNA-sequencing data

All human endometrial scRNA-seq and single-nucleus RNA-sequencing (snRNA-seq) samples were from publicly available datasets. *Fastq* files from either GEO or ArrayExpress were downloaded. Downloaded datasets included: (1) Foley *et al.* (GEO accession number, GSE260683)^28^, (2) Garcia-Alonso *et al.,* (ArrayExpress accession number, E-MTAB-10287)^29^, (3) Huang *et al.* (GEO accession number, GSE214411)^30^, (4) Lai *et al.* (GEO accession number, GSE183837)^31^, (5) Wang *et al.* (GEO accession number, GSE111976)^33^. The filtered cell/nuclei by gene matrices for Marečková *et al.* (ArrayExpress accession number, E-MTAB-14039) ^32^ were also downloaded. Only samples from healthy or SEC patients were included. Samples from patients with endometriosis were excluded. All samples were aligned to the 10X Genomics’ human reference genome GRCh38-2020-A using CellRanger.

### Single-cell and single-nuclei RNA sequencing analysis

Download and alignment of scRNA-seq data, preprocessing and batch correction, clustering parameters and annotations, and differential gene expression analysis were performed as previously described ^20^ and in Supplemental Methods. All code for preprocessing can be found on GitHub (github.com/PirtzM/EarlySEC_scRNA and https://github.com/PirtzM/SEC_Cell-Cell_Interactions). Seurat was used to process integrated data. All datasets used for analysis are listed in Supplemental Table 2.

### Interspecies transcriptomic comparisons using SAMap

For all interspecies comparisons, the filtered matrices for each sample were converted to*.h5* files for transfer to python. All further interspecies comparison analysis took place in python (v3.10.8). The filtered matrix *.h5* files for each individual sample were loaded into the environment with scanpy (v1.9.1), which were then concatenated into a single data frame per species. The annotated text files including cell barcodes, Seurat-guided cell type annotations, health status, and menstrual cycle phase were used to filter the matrix data frame based on cell barcode for comparison. Self-assembling-manifold (SAM) objects were then created for each species using the sam-algorithm package (v1.0.2). Standard preprocessing steps were followed (https://github.com/atarashansky/self-assembling-manifold, last accessed 28/07/2026)^34^.

### Cell-cell interaction analysis using CellChat

Cell interaction analysis and visualization were performed using CellChat and preprocessing was done based on the CellChat Tutorial (v2.0.1, github.com/jinworks/CellChat, last accessed 14/10/2025).

Secondary analysis used stage-specific epithelial labels transferred from our previous paper ^20^. The barcodes and associated cell type label for the cells in the stage-specific epithelial subset were transferred to the fully integrated object by matching barcodes and the luminal epithelial subclusters were regrouped into a single “LE” cluster.

### Visium spatial RNA-sequencing sample preparation and alignment

All mouse Visium spatial RNA-sequencing samples used in this manuscript were from previously reported transcriptomes^20^. As such, library preparation has been previously reported, and the previously reported sequencing data were deposited in the Gene Expression Omnibus (GEO) under the accession code GSE269332.

### Spatial RNA-sequencing spot deconvolution with *BayesPrism,* preprocessing, and analysis

Spots for each spatial RNA-sequencing sample were deconvolved using a modified version of BayesPrism^26^. Each spatial sample was deconvolved using its respective SEC stage-specific object (ie: the dysplastic single-cell object was used as a reference for all dysplastic spatial samples) from our previous paper ^20^. The output for each sample was a matrix of barcodes and cell types with a related theta value which were then added as assays to each Visium object. All preprocessing steps were performed as described in Supplemental Methods.

LE neighborhoods were labeled using the DR.SC package in R. After the adjacency matrix was created, a list of barcodes meeting specified LE criterion were labeled as “LE” based on an LE theta value greater than 0.25*sample maximum LE theta value and *Tacstd2* expression greater than 0.6*sample maximum *Tacstd2* log-normalized expression. The matrix was then filtered to only keep rows sharing a barcode with those generated in the LE list. In the resulting matrix, any barcode in rows were labeled as “LE” and any in columns were labeled as “LE_neighbor”, all other barcodes in the sample were labeled as “other”. These were added as a metadata column in each Seurat object under “neighborhood_Tacstd2”. Visual inspection of the alignment of “LE” labeled spots to the luminal structure was done and samples that included LE spots clearly unassociated with the lumen were loaded into LoupeBrowser to relabel spots for accurate representation of the epithelium.

### NOTCH2 overexpression vector cloning, transformation, and plasmid production

Mouse knock-in *Notch2* intracellular domain (N2-ICD) plasmid was created by combining N2-ICD region with a TRE promotor and an mCherry tag. The N2-ICD region was isolated and amplified from the 3XFlagN2ICD plasmid from Addgene (Addgene plasmid # 20184; kind gift from Raphael Kopan)^65^. N2-ICD was then cloned into the TR3G promoter backbone with mCherry (pTRE3G-BI-mCherry; Clontech plasmid # 631333). The new plasmid would be named TRE-N2-ICD-mCherry. All primers used for cloning can be found in Supplemental Table 3.

### Lentiviral production

A previously developed plasmid with rtTA tagged with EGFP (Addgene plasmid # 19780) was used alongside the newly cloned TRE-N2-ICD-mCherry plasmid from above. TRE-N2ICD-mCherry and rtTA-GFP lentivirus was produced in 293T cells as previously described^66^. 293T cells were seeded on 10 cm dishes at 6×10^6^ cells per dish and were verified to be attached to the dish before transfection. The TRE-N2ICD-mCherry and rtTA-GFP plasmids and PMD.2G (Addgene #12259) and PSPAX2 (Addgene #12260) were transfected using Transit-LT1 transfection reagent (Mirus bio, MIR2306) via manufacturer’s instructions. Virus was collected at 48 hours after transfection and concentrated using Lenti-X concentrator (Takara, 631231). Virus was then resuspended in organoid media with polybrene (1:1000 concentration; Sigma, TR-1003-G) and frozen at −80°C until use.

### Primary mouse endometrial organoid preparation

Three *Trp53^loxP/loxP^Rb1^loxP/loxP^* female mice were sacrificed in the diestrus stage using standard protocol. Uterine horns were collected with care to avoid uterine tube or cervical transition areas and placed in sterile 1× PBS containing 100 IU ml^-1^ of penicillin and 100 μl ml-1 streptomycin (Phosphate buffered saline pH 7.4; Corning, Corning, NY, USA, 30-002-Cl) on ice. They were then transferred to another dish of 1% Penicillin/Streptomycin in PBS on ice to wash and remove fat. Horns were then opened laterally in 4.5 mL of Collagenase/Dispase (4 µg/ml Roche Collagenase/Dispase, Sigma, Burlington, MA, 11097113001; DMEM Ham’s F12, Corning 10-092-CV) with 1 mg/mL of DNase1 (Roche, Sigma, 910104159001). The endometrium was scraped from the muscle, and all tissue was minced to about 1 mm. Minced tissue was digested in the Collagenase/Dispase solution for 45 min in the 5% CO2 incubator at 37°C. After 45 min, cell suspension was transferred to a 15 mL conical vial and mixed rigorously with a P1000. Remaining tissue pieces were left to settle and cell-containing supernatant was transferred to a new tube and rescued with 10% FBS in DMEM rescue media and placed on ice (Fraction #1). Another 4.5 mL of Collagenase/Dispase with DNaseI and another 30 min digestion step took place in the 37°C water bath for a 30 min digestion (Fraction #2). After digestion and rescue, cells were filtered into a 50 mL conical vial through a 40 μm cell strainer (Corning, 431750). Cells were embedded as a ring assay at 100,000 cells per 100 μL of Matrigel onto a 24-well cell culture-treated plate. Organoid media with ROCKi was added and plate was returned to the incubator. After 2 days of growth, ROCKi was removed from the media composition for all remaining media changes.

### NOTCH2 overexpression in endometrial organoids

Organoids were grown in Matrigel for five days. All wells were then collected using organoid harvesting media (OHM; biotechne, Minneapolis, MN 3700-100-01) into a 50 mL conical vial. Recommended steps for Matrigel dissociation were followed (biotechne Cultrex Organoid Harvesting Solution, Organoid Harvesting Procedure; https://resources.rndsystems.com/pdfs/datasheets/3700-100-01.pdf). Once all Matrigel was gone, organoids were repelleted for digestion to a single-cell level. Supernatant was removed and 2 mL of 0.25% Trypsin-EDTA (Corning, 45000-664) was added to the cells, which were placed in a 37°C water bath for a total of 15 minutes. Cells were then rescued with rescue media, spun down, resuspended in 1 mL of organoid media with ROCKi. Infections took place as described in Supplemental Methods.

After infection, cells were resuspended in Matrigel for seeding. After polymerization, organoid media + ROCKi was added to each well. Half of the wells per condition were also treated with Doxycycline (2 μg/mL; Sigma D5207). After 2 days, ROCKi was removed from the media. Doxycycline was continually provided to the cells over 9 days of growth.

### Validation of NOTCH2 overexpression in mouse endometrial organoids

The same process as above was used for NOTCH2 overexpression validation, however, wild-type C57BL6 mouse-derived endometrial organoids were used.

On day 7 and 10 after infection, organoids were imaged. On day 11, they were released from the Matrigel and then lysed for RT-qPCR using the RNA Lysis Buffer from the Zymo Quick-RNA MicroPrep kit (Zymo Research, Irvine, CA, R1050).

### RT-qPCR validation

After lysis, the standard RNA purification protocol outlined by the Zymo Quick-RNA MicroPrep kit was followed. Purified RNA was then used to make cDNA using BioRad’s iScript cDNA Synthesis kit (BioRad, Hercules, CA 1708891) in a BioRad C1000 Thermal Cycler and the recommended thermocycler settings. The RT-qPCR mix was then prepared with each condition’s cDNA, BioRad’s SsoAdvanced Universal SYBR® Green Supermix (BioRad, 1725270), and analyzed on a BioRad Thermal Cycler, Real-Time System (Bio-Rad, C1000 Touch with CFX96 Optics Module). Primer sequences are available in Supplemental Table 3.

### Endometrial organoid imaging and analysis

Organoids were manually counted under the microscope in brightfield and live-imaged using Zeiss Live-Cell Incubation Fluorescent Microscope (Zeiss Axiovert 200 M, inverted microscope). Imaging experiments were controlled using Zeiss AxioVision software. Organoids were imaged at 5X magnification in z-stacks in the brightfield, Texas red, and GFP channels. Images were flattened, processed, and analyzed using Fiji/ImageJ software (National Institutes of Health (NIH), Bethesda, MD, USA).

### Statistical analysis

Statistical comparisons were performed using a two-way ANOVA with Tukey’s multiples tests, Two-way ANOVA with Šídák’s multiple comparison, or a Kruskul-Wallis with Dunn’s multiples test in GraphPad Prism 11 software (GraphPad Software Inc., La Jolla, CA, USA). Statistical details of experiments can be found in Figure legends. Statistical significance was defined as \**P* < 0.05, \*\**P* < 0.01, \*\*\**P* < 0.001, and \*\*\*\**P* < 0.0001. Pooled data are represented as mean ± SD.

## Study approval

The Cornell University Institutional Animal Care and Use Committee (IACUC) approved all animal protocols, and experiments were performed in compliance with its institutional guidelines. De-identified sectioned slides of the endometrium of patients with and without endometrial carcinoma were prepared at Weill Cornell Medicine (IRB23-07026303) and the Johns Hopkins School of Medicine (IRB00127046). Tissue microarrays of endometrial carcinoma and normal endometrial tissue were obtained from Quickarrays Inc (Fairfield, CA, USA). These samples represent de-identified archive specimens collected by providers not involved in our research. As such, they do not meet definitions of “human participant research” under US federal regulations and Cornell IRB rules.

## Data and Code Availability Statement

The single-cell RNA-sequencing and Visium RNA-sequencing data reported in this paper are available in Gene Expression Omnibus (GEO); accession number GSE269332. All human sc/snRNA-seq samples are available from their respective authors. The Seurat objects for all analyses will be available via download on Dryad upon publication. All code for data preprocessing and Figure generation will be made available through GitHub upon publication. Any additional data supporting the findings of this study are available from the corresponding author upon reasonable request.

## Author Contributions

MGP, AFN, BDC, and AYN designed experiments. MGP, AFN, and SC performed experiments. MGP and DJP designed vectors and completed plasmid cloning. MGP, CQR, and BDC carried out bioinformatics analyses. NW, TC, and CGD developed updated BayesPrism package and completed deconvolution analysis. YZ, AY, and IMS provided human histological materials. ES, AY, IMS, and AYN performed pathological evaluations. MGP, AFN, and AYN wrote the paper.

## Acknowledgments

We thank Peter A. Schweitzer, previous Director of the Cornell Genomics Facility, Jen Greiner, current Director of the Cornell Genomics Facility, and Ann Tate, Project Manager of Transcriptional Regulation and Expression facility for their in valuable assistance with single-cell RNA-sequencing and Visium spatial transcriptomics, and Md Mozammal Hossain, Christopher S. Ashe, Caitlyn Melaram, Evelyn Kim, and Eric Lim for their excellent technical support. We also thank David McKellar and Lauren Walter for their code availability and assistance with learning methods for sequencing analysis in R. We also thank Julie Rodor and Andrew H. Baker at the University of Edinburgh for their time and expertise by sharing their Ai9-tdTomato sequence and annotation to incorporate into our single-cell and Visium spatial RNA-sequencing reference genome.

