## Supplemental Information for "Cross-species analysis links cell-cell communication rewiring to NOTCH2 during serous endometrial carcinogenesis" (from another version of this preprint: version 1, 2026-09-02)

MG Pirtz et al.

### Supplemental Methods

#### Human tissue acquisition

De-identified sectioned slides of the endometrium of patients with or without SEC, SEIC, or EEC were collected from women aged 42 - 86 years and prepared for histology at Weill Cornell Medicine and the Johns Hopkins School of Medicine. Tissue microarrays of endometrial carcinoma and normal endometrial tissue, with grade data, were obtained from Quickarrays Inc (Fairfield, CA, USA; Cat No. EMC1021).

#### Experimental animals

B6.Cg-Tg(Pax8-rtTA2S*M2)1Koes/J (Pax8-rtTA mice; RRID:IMSR_JAX:007176), B6.Cg-Tg(tetO-Cre)1Jaw/J (Tre-Cre mice; RRID:IMSR_JAX:006234) and *B6.Cg-Gt(ROSA)26Sor*$^{tm9(CAG\text{-}tdTomato)Hze}$ (Ai9 mice; RRID:IMSR_JAX:007909) were obtained from The Jackson Laboratory (Bar Harbor, ME, USA). *Trp53*$^{loxP/loxP}$ and *Rb1*$^{loxP/loxP}$ mice, which have *loxP* alleles flanking their respective genes were a gift from Dr. Anton Berns (The Netherlands Cancer Institute, Amsterdam, The Netherlands). For all experiments, mice were collected in late diestrus/early proestrus[1-4]. Estrous stage was evaluated with vaginal smears, followed by morphological verification and estimation of Ki67 index in paraffin sections of the uterus. Criteria for late diestrus stage included observing almost exclusively leukocytes in vaginal smears, a medium-wide lumen, dense/early edematous stroma, and medium-high proliferation of the glandular and luminal epithelia (Ki67 index 10–95%).

#### Doxycycline induction

Doxycycline was administered at a dose of 12 μl $g^{-1}$ body weight at a concentration of 6.7 mg $ml^{-1}$ in sterile PBS via a single intraperitoneal injection to Pax8-rtTA Tre-Cre *Trp53*$^{loxP/loxP}$*Rb1*$^{loxP/loxP}$ Ai9 mice and control mice at 6–24 weeks old, as previously described[5,6]. Mice were identified to be in late diestrus phase before induction using vaginal smears. All mice were euthanized by $CO_2$ and further analyses were carried out.

**Pathological evaluation**

All mice underwent gross pathology evaluation at the time of necropsy, followed by histology, immunohistochemistry, and image analysis. Potential sites of endometrial carcinoma metastasis were evaluated carefully, including the local lymph nodes, omentum, peritoneum, and lungs. Dysplastic samples were diagnosed based on atypical nuclear to cytoplasmic ratios, mitotic figures, and atypical glandular patterning in the endometrial epithelium, as previously described[5]. Pre-dysplastic samples lacked morphologically discernable neoplastic lesions but were marked by presence of tdTomato/RFP staining indicative of Cre-*LoxP* mediated recombination. All human cases underwent standardized pathology review and H-score semi-quantification.

**Histology, immunohistochemistry, and image analysis**

All tissues were fixed in buffered 4% paraformaldehyde overnight at 4°C followed by standard tissue processing and paraffin embedding. Histology and immunohistochemistry staining was carried out on 4-μm-thick tissue sections. For immunohistochemistry, antigen retrieval was performed as needed by incubation of deparaffinized and rehydrated tissue sections in boiling 10 mM sodium citrate buffer (pH 6.0) for 15 min unless otherwise noted. The primary antibodies were incubated at 4°C overnight or 1.5 h at room temperature (RT), followed by incubation with secondary biotinylated or fluorophore conjugated antibodies (1 h, RT). Incubation with biotinylated antibodies slides followed by modified Elite avidin-biotin peroxidase (ABC) method (Vector Laboratories, Burlingame, CA, USA; pk-6100) for 30 min (RT). All antibodies used for immunostaining are listed in Supplemental Table 1. Stained sections were scanned by ScanScope CS2 (Leica Biosystems, Vista, CA, USA) with a 40× objective, followed by the analysis with Fiji/ImageJ software (National Institutes of Health (NIH), Bethesda, MD, USA). Following Vectashield Vibrance Antifade Mounting Medium with DAPI (H-1800, Vector Laboratories), immunofluorescence samples were tilescanned (Zeiss 710 upright Confocal, Cornell University's Biotechnology Resource Center) at 40×. Analysis was performed with Fiji/ImageJ software (NIH).

**Single-cell RNA-sequencing library preparation**

All mouse single-cell RNA-sequencing (scRNA-seq) samples used in this manuscript were from previously reported transcriptomes[5]. As such, library preparation has been previously reported and data deposited in the Gene Expression Omnibus (GEO) under the accession code GSE269332.

**Download and alignment of single-cell and single-nucleus RNA-sequencing data**

All mouse samples were previously published and can be accessed from GSE269332. Alignment of these samples were previously described[5].

All human endometrial scRNA-seq and single-nucleus RNA-sequencing (snRNA-seq) samples were from publicly available datasets. *Fastq* files from either GEO or ArrayExpress were downloaded. Downloaded datasets included: (1) Foley *et al.* (GEO accession number, GSE260683)[7], (2) Garcia-Alonso *et al.,* (ArrayExpress accession number, E-MTAB-10287)[8], (3) Huang *et al.* (GEO accession number, GSE214411)[9], (4) Lai *et al.* (GEO accession number, GSE183837)[10], (5) Wang *et al.* (GEO accession number, GSE111976)[11]. The filtered cell/nuclei by gene matrices for Marečková *et al.* (ArrayExpress accession number, E-MTAB-14039)[12] were also downloaded. Only samples from healthy or SEC patients were included. Samples from patients with endometriosis were excluded.

Since raw reads for the Marečková *et al.* dataset were unavailable, we aligned all other samples to the same reference. All samples were aligned to the 10X Genomics' human reference genome GRCh38-2020-A using the CellRanger (v7.1.0, 10X Genomics) *count* function.

**Mouse single-cell RNA-sequencing preprocessing and batch correction**

All preprocessing of mouse scRNA-seq data was done as previously described[5]. All code for preprocessing can be found on GitHub (github.com/PirtzM/EarlySEC_scRNA). Seurat was used to process integrated data.

**Mouse dataset clustering parameters and annotations**

Original dataset preparation was done as previously described[5]. The fully integrated dataset was used for all mouse scRNA-seq analysis in this manuscript. Cell clustering was determined using Louvain clustering on the output graphs (*Seurat::FindClusters*). Cell types were determined based on commonly used genes for the healthy, adult mouse uterus, and as previously described[13,14]. References from the mouse uterus at diestrus were used when possible.

Individual SEC stage-specific datasets and epithelial subpopulations were prepared for Visium spot deconvolution and higher resolution CellChat, as previously described[5].

**Human single-cell RNA-sequencing preprocessing and batch correction**

Since the unfiltered matrices for Marečková *et al.* datasets were unavailable, ambient RNA removal was unable to be done, so this step was skipped. The standard Seurat (v4.3.0) workflow (*NormalizeData*, *ScaleData*, *FindVariableFeatures*, *RunPCA*, *FindNeighbors*, *FindClusters*, and *runUMAP*; github.com/satijalab/seurat) was used for Seurat object preparation and preprocessing. DoubletFinder (v2.0.3;github.com/chris-mcginnis-ucsf/DoubletFinder, last accessed 06/08/2025) was used to predict doublets in each dataset individually. The default $P_N$ value (0.25) was used, and $BC_{mvn}$ optimization was used to identify ideal $P_K$ values for each sample. Doublet rate was informed by the 10X Chromium Handbook. Then, all datasets were merged. Cells with fewer than 200 features (nFeature), fewer than 750 transcripts (nCounts), or more than 25% of unique transcripts related to mitochondrial genes, and predicted doublets were removed. Samples SE02_A, SE02_B, SE03_A, SE03_B, and P3 were eliminated based on QC parameters and doublet predictions. After preprocessing and merging of the datasets, batch correction was performed using Harmony (v0.1.1;

github.com/immunogenomics/harmony, last accessed 06/08/2025). Before final analyses and cluster naming, deceased samples and samples with non-standard menstrual phase labels (ie: “5 days after ovulation”, “Unknown”, and “Hormones”) were removed (*Seurat::subset*) to avoid confounding gene expression patterns. The dataset was rescaled (*Seurat::FindVariableFeatures* and *Seurat::ScaleData*), reintegrated, reharmonized, and reclustered, then Seurat was used to process the integrated data. This dataset will be further referred to as the “All Menstrual Phase dataset”.

**Human dataset clustering parameters and annotations**

Cell clustering was determined using Louvain clustering on the output graphs (*Seurat::FindClusters*). Clustering resolution and K values were adjusted for each combination of samples to best represent the data. Cell types were determined based on commonly used genes for the healthy, cycling human uterus, as previously described[8,11,12].

The initial human dataset (“All Menstrual Phase dataset”) used in SAMap to determine correlation between human menstrual cycle phase and normal mouse diestrus phase included epithelial and cycling cells subset (*Seurat::subset*) from the original data object. Cells were broadly clustered based on canonical epithelial (*EPCAM, KRT8, PAX8*), stromal (*PDGFRA, DIO2, PECAM1, ACTA2, PTPRC*), and cycling (*MKI67*) gene expression.

Once the proliferative early and proliferative phases were chosen (to be labeled as “Final Human Dataset”), these samples, alongside the SEC samples, were subset (*Seurat::subset),* rescaled (*Seurat::FindVariableFeatures* and *Seurat::ScaleData*), reintegrated, reharmonized, and reclustered. Clusters were named as described above. Notably, some of our epithelial populations express immune-associated genes. It is possible some cells were contaminated with ambient immune cell RNA, as ambient RNA removal was unable to be done. An additional cell type metadata column was added to the object which grouped all similar epithelial populations together to leave only a singular cluster of each LE, GE, Hormone-responsive Progenitor-like, Ciliated, and Cycling. This metadata column was labeled “seurat_clusters_CC”.

All mouse and human datasets used throughout the manuscript are described in Supplemental Table 2.

**Interspecies transcriptomic comparisons using SAMap**

For robust comparisons between mouse and human samples, we aimed to find the human menstrual cycle phase most similar to the normal, diestrus mouse uterus. The epithelial and cycling cells from the mouse dataset and “All Menstrual Phase” human dataset were subset (*Seurat::subset*). An annotated text file containing barcodes, menstrual phase (human) or SEC stage (mouse), and Seurat-guided cell type annotations was created.

The filtered matrices for each sample were also converted to *.h5* files (*scCustomize::Create_10X_H5*; v1.1.3) for transfer to python. All further interspecies comparison analysis took place in python (v3.10.8). The filtered matrix *.h5* files for each individual sample were loaded into the environment with scanpy (v1.9.1), which were then concatenated into a single data frame per species. The annotated text files were used to filter the matrix data frame based on cell barcode to keep only the epithelial and cycling cells. Self-assembling-manifold (SAM) objects were then created for each species using the sam-algorithm package (v1.0.2). Standard preprocessing steps were followed (https://github.com/atarashansky/self-assembling-manifold, last accessed 28/07/2026)[15]. Once species-specific SAM objects were created, the SAMap (v 1.0.12) pipeline was applied to the SAM objects to map the mouse and human datasets together.

This workflow was repeated once early proliferative, proliferative, and SEC samples were subset ("Final Human Dataset") from the "All Menstrual Phase" human dataset to gain a better understanding of how Normal and SEC cell types relate between the two species. The epithelial and cycling cells were subset in R (*Seurat::subset*) from the "Final Human Dataset" and an annotated text file containing cell barcodes, associated health status (normal or SEC), and Seurat-guided cell type annotations was created. This was repeated for the mouse dataset.

The two subsets (human epithelial/cycling and mouse epithelial/cycling) were further split into healthy and SEC groups and then subjected to the same protocol as above in python to create four SAM objects. The healthy human and mouse epithelial/cycling SAM objects and the SEC human and mouse epithelial/cycling SAM objects were mapped based on cluster names from Seurat using the SAMap pipeline for cell type correlation analysis and gene-gene mapping between species.

**Cell-cell interaction analysis using CellChat**

Cell interaction analysis and visualization were performed using CellChat (v2.0.1, github.com/jinworks/CellChat, last accessed 14/10/2025). CellChat preprocessing of the mouse scRNA-seq Seurat object was done based on the CellChat Tutorial (v2.0.1, github.com/jinworks/CellChat, last accessed 14/10/2025). The fully integrated object was split by SEC stage using *Seurat::SplitObject()*. Each SEC stage object was then turned into a CellChat object using *CellChat::createCellChat()*, and each stage was processed individually following the CellChat tutorial pipeline and the mouse signaling database (*CellChatDB.mouse*). Key changes from the tutorial include setting "raw.use=TRUE" and "population.size=TRUE" in *CellChat::computeCommunProb()* step.

For higher resolution analysis, stage-specific epithelial labels were added from our previous paper[5]. The barcodes and associated cell type label for the cells in the stage-specific epithelial subset were extracted from their objects. These labels were transferred to the fully integrated object by matching barcodes using the *hash::hash()* function (v2.2.6.3) and the luminal epithelial subclusters were regrouped into a single "LE" cluster. Any remaining cells in the fully integrated epithelial bins without associated stage-specific epithelial barcodes were renamed based on their cell type from the All cell, stage-specific datasets. These cells were likely misclustered based on disease-associated cell changes. The new metadata column was labeled as "seurat_clusters_mergeGroup". With these new labels, new CellChat objects were built with the same parameters as above but now grouped by the "seurat_clusters_mergeGroup" column.

**Visium spatial RNA-sequencing sample preparation and alignment**

All mouse Visium spatial RNA-sequencing samples used in this manuscript were from previously reported transcriptomes[5]. As such, library preparation has been previously reported, and the previously reported sequencing data were deposited in the Gene Expression Omnibus (GEO) under the accession code GSE269332.

**Spatial RNA-sequencing spot deconvolution with *BayesPrism***

Spots for each spatial RNA-sequencing sample were deconvolved using a modified version of BayesPrism[16] (https://github.com/Danko-Lab/BayesPrism), which uses a similar Bayesian prediction model to deconvolve spatial RNA-sequencing spots as BayesPrism uses for bulk RNA-sequencing datasets but can also receive multiple references for annotation.

Alongside the count matrices from the Visium spatial RNA-sequencing samples, the SEC stage-specific scRNA-seq objects were loaded in as references. The scRNA-seq object provided the average gene expression profile of each annotated cell type. PrismST then used this information to predict the cell type composition of each spatial spot barcode through *PrismST::run_prismST()* for each Visium sample. The output for each sample was a matrix of barcodes and cell types with a related theta value representing the estimated proportion of each cell type in the barcode.

Each spatial sample was deconvolved using its respective SEC stage-specific object (ie: the dysplastic single-cell object was used as a reference for all dysplastic spatial samples). First, the SEC-stage specific integrated single-cell dataset with both epithelial and stromal cells from our previous paper[5], was used as a reference to deconvolve each spot barcode. Then, the SEC stage-specific epithelial subsets were used for a higher epithelial cell type resolution deconvolution of the spot barcodes.

**Spatial RNA-sequencing preprocessing**

Filtered feature matrices from the aligned files were loaded into R using *Seurat::Load10X_Spatial*. Metadata features including percent mitochondrial reads, hemoglobin scores, and ribosomal protein scores were added. Spots with fewer than 1,000 reads and 500 UMIs were removed with the assumption they were low-quality spots.

Two assays were added to each spatial Seurat object from the BayesPrism theta outputs. The first, "predictions", included the theta values for all cell types as rows and barcodes as the columns. The second, "predictions_merge", included broad cell type identifiers, which were calculated using the summed theta values for similar cell types (ie: "Fib 1" and "Fib 2" became "Fib"). Most analyses requiring cell type labels were performed using the "predictions_merge" assay.

Finally, since QC occurred after BayesPrism, the predictions assays included a few extra cells. All low-quality spot barcodes were removed from these assays to match the metadata and Spatial assays.

**Luminal neighborhood analysis of spatial RNA-sequencing data**

The spot coordinates were added to the metadata of each spatial Seurat object for easy access in later steps. Using the DR.SC package in R (v3.4; https://github.com/feiyoung/DR.SC, last accessed 14/10/2025), an adjacency matrix was created for each sample (*DR.SC::getAdj()*), where the hexagonal grid characteristic of Visium was used to assume each spot had six adjacent spots. The matrix was populated with a 1 if spots were adjacent to one another and a 0 if they were not and barcodes were added back to the matrix. A list of barcodes meeting specified LE criterion were labeled as “LE” based on an LE theta value greater than 0.25*sample maximum LE theta value and *Tacstd2* expression greater than 0.6*sample maximum *Tacstd2* log-normalized expression.

The matrix was then filtered to only keep rows sharing a barcode with those generated in the LE list. The columns containing zeros were removed. In the resulting matrix, any barcode in rows were labeled as “LE” and any in columns were labeled as “LE_neighbor”, all other barcodes in the sample were labeled as “other”. These were added as a metadata column in each Seurat object under “neighborhood_Tacstd2”. For more robust feature plotting, this column was also duplicated as pseudo-continuous variables, where “LE” labels were replaced with a 10, “LE_neighbor” labels were replaced with a 5, and all “other” labels were replaced with a 0. These were saved under “neighborhood_5”.

Visual inspection of the alignment of “LE” labeled spots to the luminal structure was done. Samples that included LE spots clearly unassociated with the lumen were loaded into LoupeBrowser (v7.0.1, 10X Genomics), and the spots were manually relabeled. The resulting barcode by LE neighborhood matrix was then exported from LoupeBrowser and reloaded into R to update the “neighborhood_Tacstd2” metadata column.

**NOTCH2 overexpression vector cloning, transformation, and plasmid production**

Mouse knock-in *Notch2* intracellular domain (N2-ICD) plasmid was created by combining N2-ICD region with a TRE promotor and an mCherry tag. The N2-ICD region was isolated and amplified from the 3XFlagN2ICD plasmid from Addgene (Addgene plasmid # 20184; kind gift from Raphael Kopan)[17]. N2-ICD was then cloned into the TR3G promoter backbone with mCherry (pTRE3G-BI-mCherry; Clontech plasmid # 631333). The new plasmid would be named TRE-N2-ICD-mCherry. All primers used for cloning can be found in Supplemental Table 3.

**Lentiviral production**

A previously developed plasmid with rtTA tagged with EGFP (Addgene plasmid # 19780) was used alongside the newly cloned TRE-N2-ICD-mCherry plasmid from above. TRE-N2ICD-mCherry and rtTA-GFP lentivirus was produced in 293T cells as previously described[18]. 293T cells were seeded on 10 cm dishes at $6x10^6$ cells per dish and were verified to be attached to the dish before transfection. The TRE-N2ICD-mCherry and rtTA-GFP plasmids and PMD.2G (Addgene #12259) and PSPAX2 (Addgene #12260) were transfected using Transit-LT1 transfection reagent (Mirus bio, MIR2306) via manufacturer's instructions. Virus was collected at 48 hours after transfection and concentrated using Lenti-X concentrator (Takara, 631231). Virus isolated by centrifuging at max speed at 4°C for 45 minutes. Virus was then resuspended in organoid media with polybrene (1:1000 concentration; Sigma, TR-1003-G) and frozen at -80°C until use.

**Primary mouse endometrial organoid preparation**

Three *Trp53*$^{loxP/loxP}$*Rb1*$^{loxP/loxP}$ female mice were sacrificed in the diestrus stage using standard protocol. Uterine horns were collected with care to avoid uterine tube or cervical transition areas and placed in sterile 1× PBS containing 100 IU ml$^{-1}$ of penicillin and 100 μl ml-1 streptomycin (Phosphate buffered saline pH 7.4; Corning, Corning, NY, USA, 30-002-CI) on ice. They were then transferred to another dish of 1% Penicillin/Streptomycin in PBS on ice to wash and remove fat. Horns were then opened laterally in 4.5 mL of Collagenase/Dispase (4 μg/ml Roche Collagenase/Dispase, Sigma, Burlington, MA, 11097113001; DMEM Ham's F12, Corning 10-092-CV) with 1 mg/mL of DNase1 (Roche, Sigma, 910104159001). The endometrium was scraped from the muscle, and all tissue was minced to about 1 mm. The dish containing minced tissue and Collagenase/Dispase solution was placed in the 5% $CO_2$ incubator at 37°C for 45 min. After 45 min, cell suspension was transferred to a 15 mL conical vial and mixed rigorously with a P1000. Remaining tissue pieces were left to settle and cell-containing supernatant was transferred to a new tube and rescued with 10% FBS in DMEM rescue media (DMEM Ham's F12, Corning 10-092-CV; 20% fetal bovine serum [FBS], Sigma-Aldrich, St. Louis, MO, USA, F4135) and placed on ice (Fraction #1). Another 4.5 mL of Collagenase/Dispase with DNaseI was added to the remaining tissue pieces and mixed again vigorously with a P1000. They were then placed in a 37°C water bath for a 30 min digestion. Every 15 minutes, the tissue was mixed with a P1000. After 30 minutes, cells were rescued with rescue media (Fraction #2). The contents of both the Fraction #1 and Fraction #2 tubes were filtered into a 50 mL conical vial through a 40 μm cell strainer (Corning, 431750). The back of a new P1000 filter tip was used to gently push cells through the filter, and then 1 mL of rescue media was used to rinse the filter. Cells were then pelleted by centrifugation at 600 x g for 5 min at 4°C. Supernatant was removed and 1 mL of rescue media was added to resuspend the cells. Cells were then counted using a hemocytometer, and the number of collected cells informed the number of wells to be seeded. Cells were embedded as a ring assay at 100,000 cells per 100 µL of Matrigel onto a 24-well cell culture-treated plate. Organoid media (Gibco Advanced DMEM/F12, Thermo Fisher 12634010; 25% L-WRN Conditioned Medium, ATCC CRL-3276; 12 mM HEPES, Thermo Fisher 15630080; 1% GlutaMax, Thermo Fisher 35050079; 2% B27, Thermo Fisher 17504044; 1% N2, Thermo Fisher 17502048; 100 ng/ml hEGF, Sigma E9644; 100 ng/ml Human FGF-10, 1 mM Nicotinamide, Sigma N0636; 2.5 µM SB431542 [TGF-B RI Kinase Inhibitor VI], Selleckchem, Houston, TX S1067) was then added with 10 µM Y-27632 (ROCKi; Sellckchem S1049) and plate was returned to the incubator. After 2 days of growth, ROCKi was removed from the media composition for all remaining media changes. Media was changed every 2-3 days.

**NOTCH2 overexpression in endometrial organoids**

Organoids were grown in Matrigel for five days. All wells were then collected using organoid harvesting media (OHM; biotechne, Minneapolis, MN 3700-100-01) into a 50 mL conical vial. Recommended steps for Matrigel dissociation were followed (biotechne Cultrex Organoid Harvesting Solution, Organoid Harvesting Procedure; https://resources.rndsystems.com/pdfs/datasheets/3700-100-01.pdf). Once all Matrigel was gone, organoids were repelleted for digestion to a single-cell level. Supernatant was removed and 2 mL of 0.25% Trypsin-EDTA (Corning, 45000-664) was added to the cells, which were then mixed vigorously 30 times. The tube was then placed in a 37°C water bath for a total of 15 minutes with breaks every 5 minutes to mix. After 15 minutes, the cells were vigorously mixed 100 times and 4 mL of rescue media was added to stop the trypsin digestion. Now at a single-cell level, the cells were spun down, supernatant removed and resuspended in 1 mL of organoid media + ROCKi.

Cells were counted and calculations were done to provide 30,000 cells per infection well in a 96-well cell culture-treated plate (Corning, 5396) across four infection conditions with two infection wells per condition. Infection conditions were as follows: AdenoCRE + TRE-N2-mCherry + rtTA-GFP, AdenoBlank + TRE-N2-mCherry + rtTA-GFP, AdenoCRE only, and AdenoBlank only. Cells were infected with Adenoviruses at an MOI of 500 and 100 µL of the N2-ICD overexpression virus (containing both the TRE-N2-mCherry and rtTA-GFP elements) was added to the infection wells. Infection wells without N2-ICD overexpression virus were supplemented with an additional 100 µL of organoid media + ROCKi. The cells were then subjected to a 1-hour spinfection at RT and then transferred to a 37°C incubator for another 2.5 h.

After infection, the supernatant from the wells were collected into individual Eppendorf tubes and placed on ice (two tubes per infection condition, four infection conditions). Then, 100 µL of 0.25% Trypsin-EDTA was placed into each well and the 96-well plate was returned to the incubator for 1 minute. Once any attached cells were released from the bottom of the well, the trypsin cell suspension was added to its corresponding Eppendorf tube and the digestion was stopped. Cells were spun down 600 x g for 5 min at 4°C and supernatant was removed.

Finally, cells were resuspended in 120 µL of Matrigel to seed twelve-20 µL droplets per condition across two 24-well cell culture-treated plates. Once droplets were placed in the wells, the plates were returned to the 37°C incubator for 1.25 h to allow for full Matrigel polymerization. At that time, 300 µL of organoid media + ROCKi was added to each well.

Half of the wells per condition were also treated with Doxycycline (2 µg/mL; Sigma D5207). After 2 days, ROCKi was removed from the media. Doxycycline was continually provided to the cells over 9 days of growth. Media was refreshed every 2 days.

**Validation of NOTCH2 overexpression in mouse endometrial organoids**

Cells from passage 13, wild-type C57BL6 endometrial organoids were thawed and added to our 10% FBS in DMEM rescue media. The cells were pelleted at 600 x g for 5 min and resuspended in 1 mL of rescue media for counting with a hemocytometer. After counting, cells were pelleted again and supernatant removed. Cells were then resuspended in Matrigel (Corning, CLS356231) to seed 100 µL rings with 100,000 cells per ring in a 24-well cell culture-treated plate (Corning, 353047). After 1 h in the 5% $CO_2$ incubator at 37°C, endometrial organoid media with ROCKi was added. Organoids were cultured for 10 days. ROCKi was removed from media after day 2.

After 10 days, organoids were passed and returned to a single-cell suspension using the same protocol as above. Now at a single-cell level, the cells were spun down, supernatant removed and resuspended in 1 mL of organoid media + ROCKi. Cells were counted and calculations were done to provide 30,000 cells per infection well in a 96-well cell culture-treated plate (Corning, 5396) across two infection conditions and a control, no infection condition: TRE-N2-mCherry + rtTA-GFP and TRE-N2-mCherry only. Once seeded, 100 µL of the N2-ICD overexpression virus (containing both the rtTA-GFP and TRE-N2-mCherry elements) and TRE-N2-mCherry virus was added to the infection wells. An additional 100 µL of organoid media + ROCKi was added to all wells to prevent cells drying out. The same infection and collection protocol as above was used. Cells were spun down 600 x g for 5 min at 4°C and supernatant was removed.

Finally, cells were resuspended in Matrigel to seed 6-wells per condition in one 20 µL drop per well. Each droplet contained 5,000 cells. Once droplets were placed in the wells, the plates were returned to the 37°C incubator for 1.25 h to allow for full Matrigel polymerization. At that time, 300 µL of organoid media + ROCKi was added to each well. Half of the wells per condition were also treated with Doxycycline (2 µg/mL; Sigma, d5207) to activate the transcription of the N2-ICD-mCherry. After 2 days, ROCKi was removed from the media. Doxycycline was continually provided to the cells over 11 days of growth. Media was refreshed every 2 days.

On day 7 and 10 of growth, organoids were imaged. On day 11, they were released from the Matrigel and then lysed for RT-qPCR using the RNA Lysis Buffer from the Zymo Quick-RNA MicroPrep kit (Zymo Research, Irvine, CA, R1050).

**RT-qPCR validation**

After lysis, the standard RNA purification protocol outlined by the Zymo Quick-RNA MicroPrep kit was followed. Purified RNA was then used to make cDNA using BioRad's iScript cDNA Synthesis kit (BioRad, Hercules, CA 1708891) in a BioRad C1000 Thermal Cycler and the recommended thermocycler settings. The RT-qPCR mix was then prepared with each condition's cDNA, BioRad's SsoAdvanced Universal SYBR® Green Supermix (BioRad, 1725270), and analyzed on a BioRad Thermal Cycler, Real-Time

System (Bio-Rad, C1000 Touch with CFX96 Optics Module). Primer sequences are available in Supplemental Table 3.

#### Endometrial organoid imaging and analysis

After nine days of growth, organoids were manually counted under the microscope in brightfield. Then, organoids were live imaged using Zeiss Live-Cell Incubation Fluorescent Microscope (Zeiss Axiovert 200 M, inverted microscope) and imaging experiments were controlled using Zeiss AxioVision software. During imaging, organoids were maintained at 37°C. Organoids were imaged at 5X magnification in z-stacks in the brightfield, Texas red, and GFP channels. Images were flattened, processed, and analyzed using Fiji/ImageJ software (National Institutes of Health (NIH), Bethesda, MD, USA).

#### Statistical analysis

Statistical comparisons were performed using a two-way ANOVA with Tukey's multiples tests, Two-way ANOVA with Šídák's multiple comparison, or a Kruskul-Wallis with Dunn's multiples test in GraphPad Prism 11 software (GraphPad Software Inc., La Jolla, CA, USA). Statistical details of experiments can be found in Figure legends. Statistical significance was defined as *$P < 0.05$, **$P < 0.01$, ***$P < 0.001$, and ****$P < 0.0001$. Pooled data are represented as mean ± SD.

#### Study approval

The Cornell University Institutional Animal Care and Use Committee (IACUC) approved all animal protocols, and experiments were performed in compliance with its institutional guidelines. De-identified sectioned slides of the endometrium of patients with or without SEC, SEIC, EEC were prepared at Weill Cornell Medicine (IRB23-07026303) and the Johns Hopkins School of Medicine (IRB00127046). Tissue microarrays of endometrial carcinoma and normal endometrial tissue were obtained from Quickarrays Inc (Fairfield, CA, USA). These samples represent de-identified archive specimens collected by providers not involved in our research. As such, they do not meet definitions of "human participant research" under US federal regulations and Cornell IRB rules.

**Supplemental Figures**

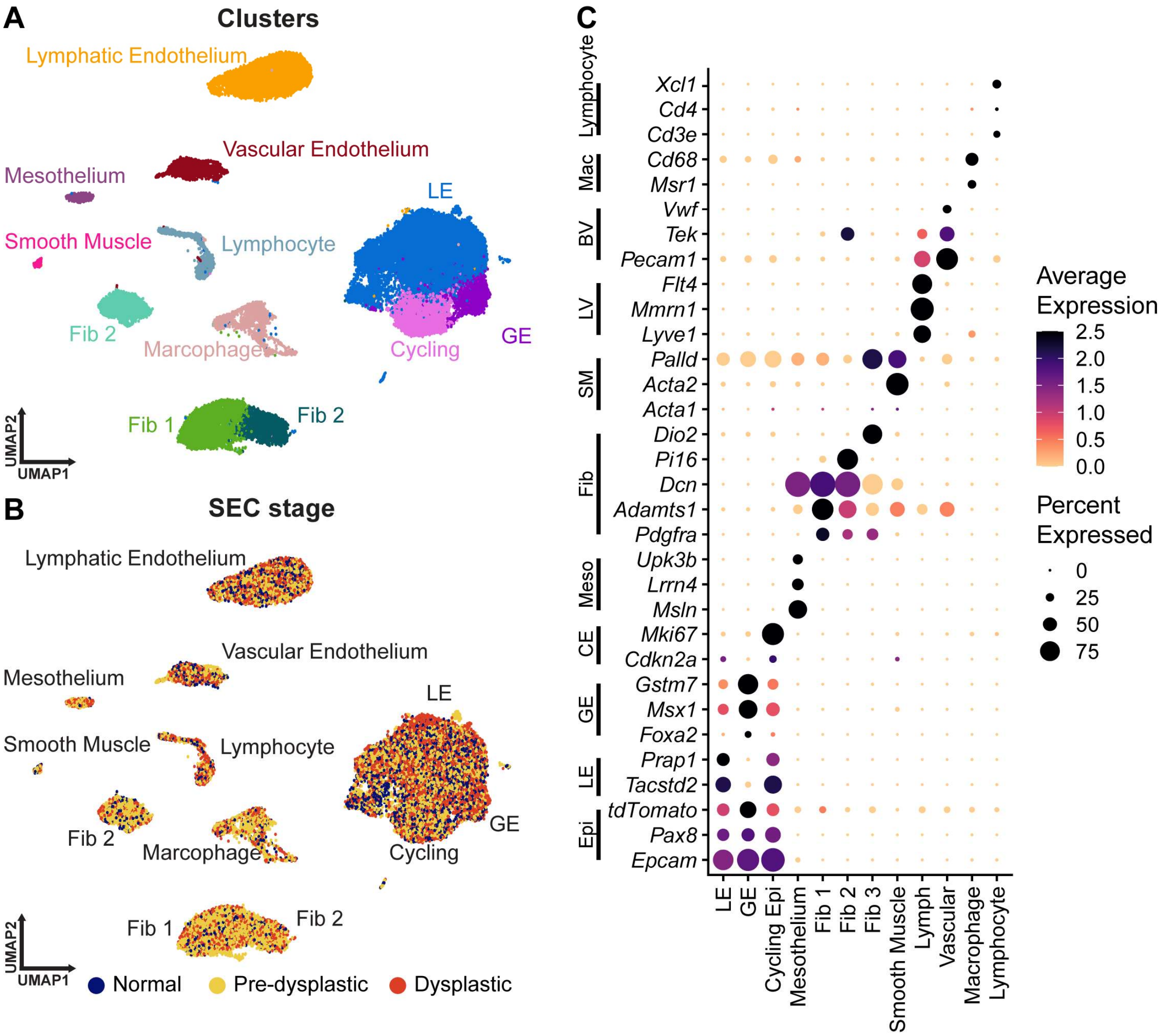

**Supplemental Figure 1. Mouse endometrial single-cell RNA sequencing atlas** (A-B) UMAP, the fully integrated dataset, colored by cell type cluster (A) and colored by SEC stage (B). (C) Average log-normalized expression of genes used to name clusters in the fully integrated dataset. LE, Luminal Epithelium; GE, Glandular Epithelium; Fib, Fibroblast; Lymph, Lymphatic Endothelium. tdTomato was aligned under the gene name "TdTomato-UTR".

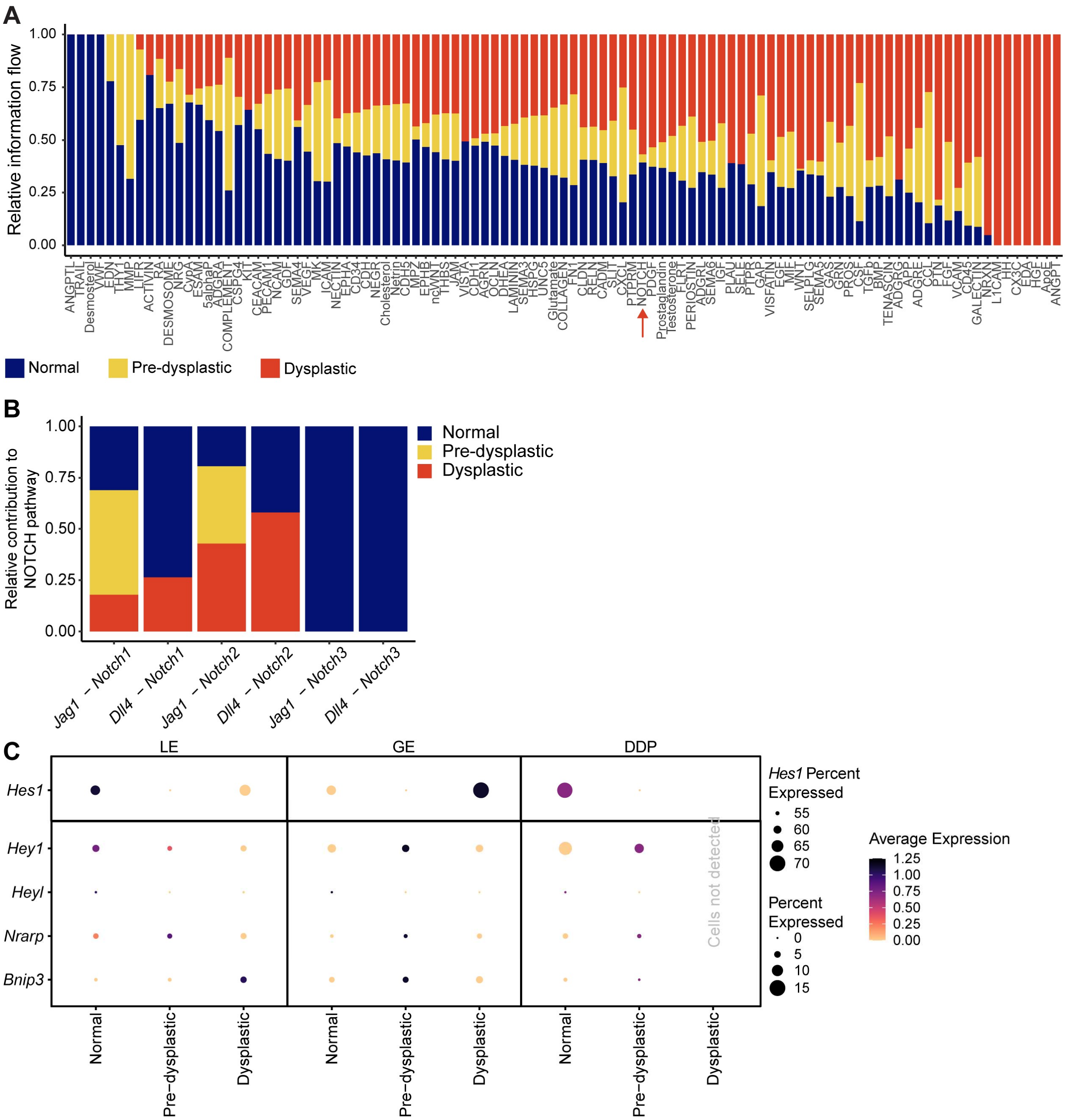

**Supplemental Figure 2. Interaction strengths and contribution trends as predicted by CellChat.** (A) The relative contribution of each SEC stage to an individual CellChat annotated pathway based on interaction strengths. Arrow, highlighting the NOTCH pathway. (B) Relative contribution of each NOTCH pathway L-R pairs to interactions involving epithelial cell receivers to each stage based on weight. (C) Average log-normalized NOTCH pathway target gene expression in each epithelial population in each SEC stage.

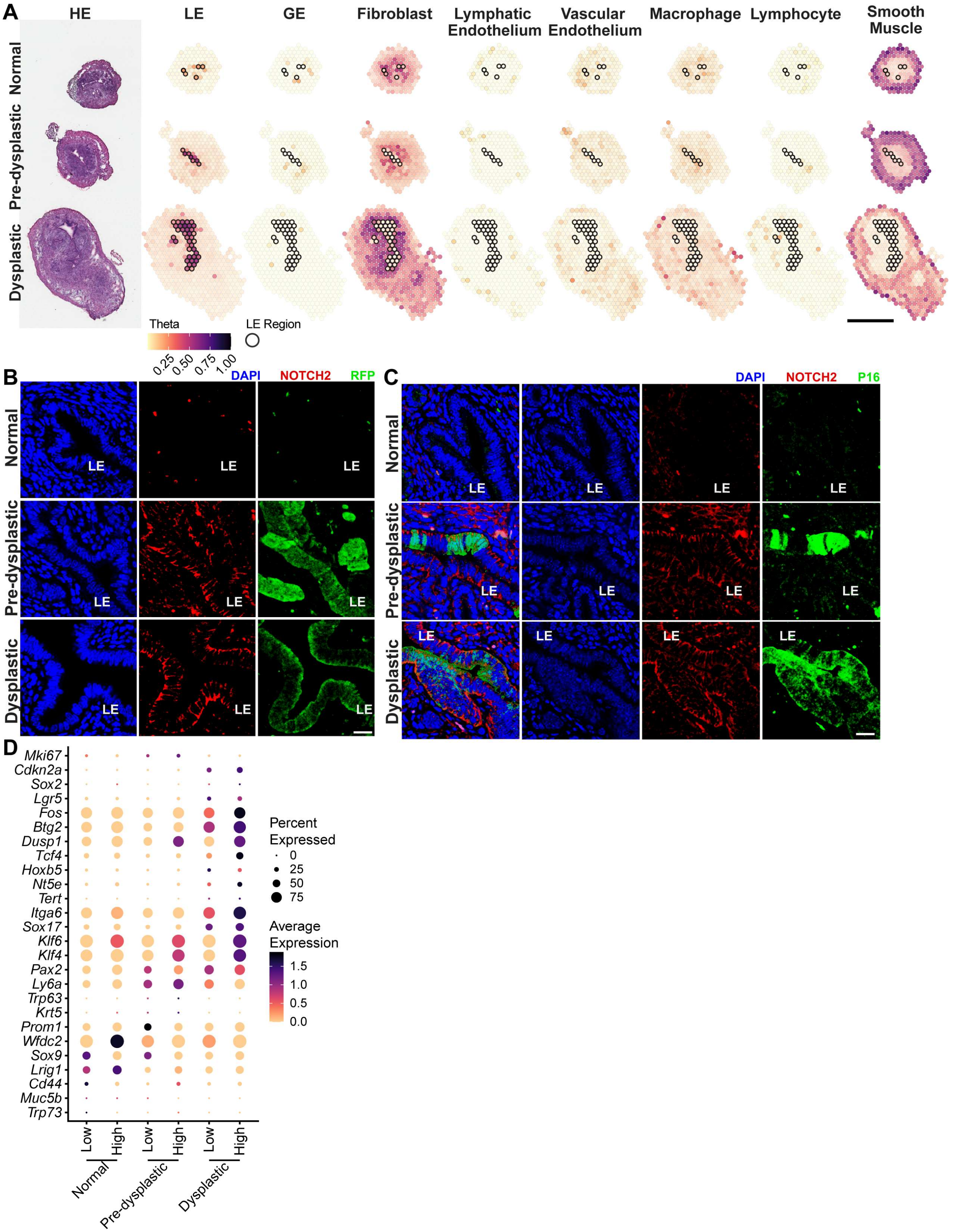

**Supplemental Figure 3. Spatial transcriptomic spot deconvolution and NOTCH pathway component expression.** (A) BayesPrism theta predictions for main cell types in the Spatial RNA-sequencing data. LE spots (LE region) marked by black outline. Scale bar, 2 mm for all images. (B-C) Representative images of NOTCH2 (red) and RFP (B; green) or P16 (C; green) in normal, pre-dysplastic, and dysplastic samples. Immunofluorescence with DAPI (blue) counterstain. Confocal microscopy. Scale bar in bottom right image, 20 µm for all images. (D) Average log-normalized stem-like gene expression in the grouped LE, GE, and DDP populations in each SEC stage, stratified by *Notch2*-expression. Low, below the average normal epithelial expression of *Notch2*; High, above the average normal epithelial expression of *Notch2*.

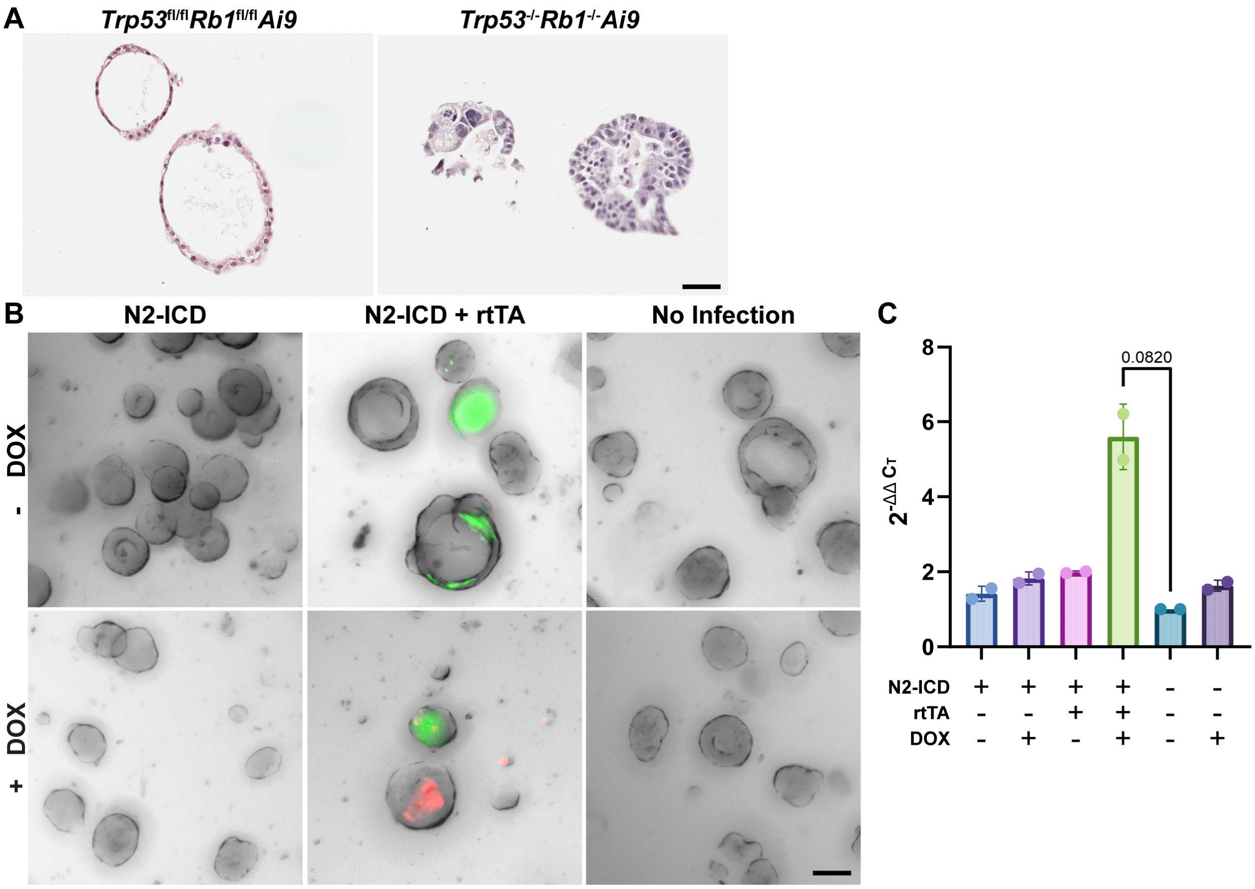

**Supplemental Figure 4. Experimental organoid system and validation of inducible NOTCH2 activation.** (A) Endometrial epithelial organoids show dysplastic morphology with cell crowding and atypical nuclei after inactivation of *Trp53* and *Rb1* (right image). Hematoxylin and eosin. Scale bar, 45 µm for both images. (B) Representative organoids 7 days after infection. Live fluorescent cell imaging. Scale bar in bottom right image, 100 µm for all images. (C) RT-qPCR detection of the *Notch2* intracellular domain in the organoids in each condition. Normalized to GAPDH expression and *Notch2* intracellular domain expression in the no infection control. N2-ICD, TRE-N2ICD-mCherry virus infection; rtTA, rtTA-GFP virus infection; DOX, Doxycycline. Each point is a RT-qPCR well replicate. Kruskul-Wallis, Dunn's multiples test. Error bars denote SD.

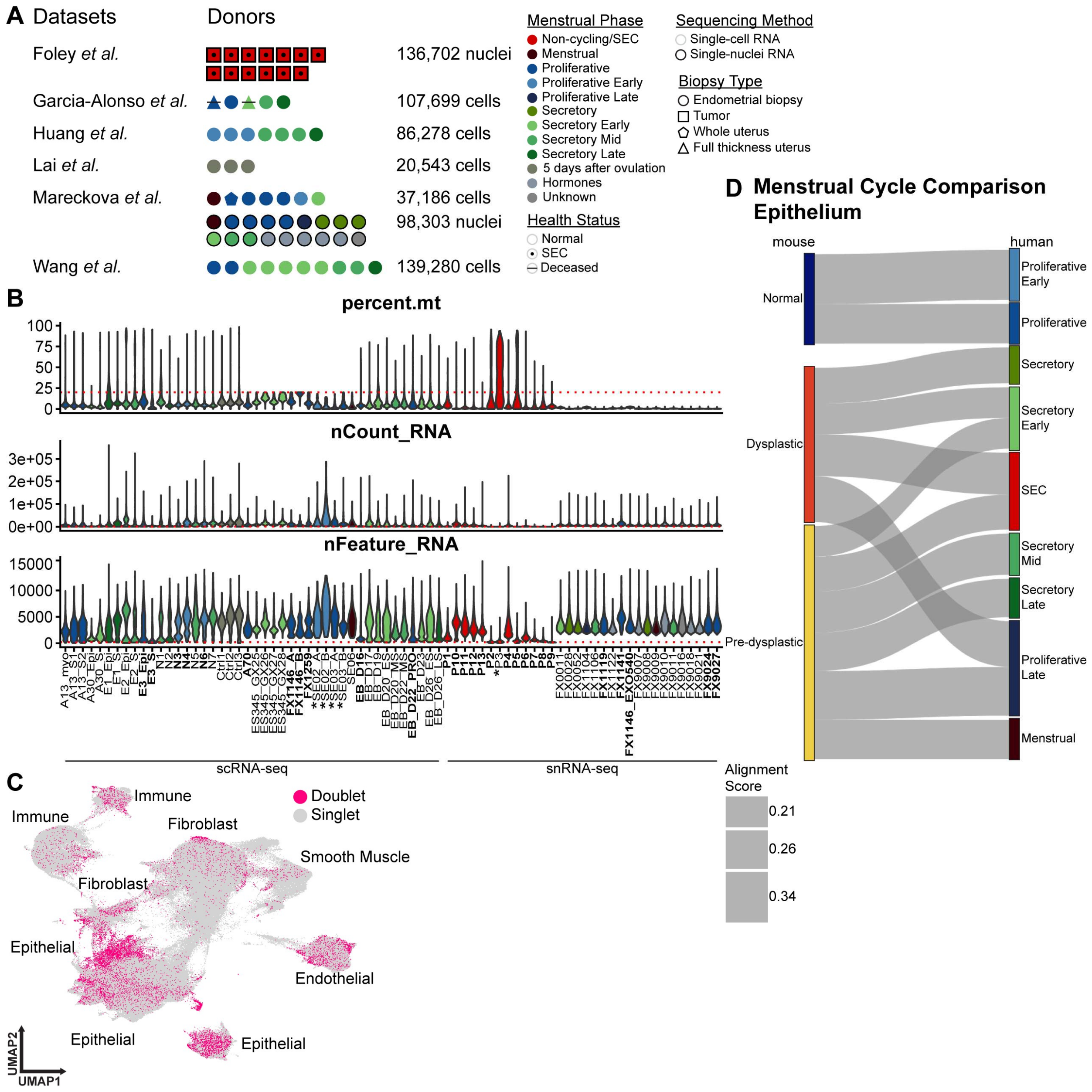

**Supplemental Figure 5. Human RNA sequencing dataset preprocessing.** (A) Schematic of samples collected from publicly available human endometrial datasets. (B) Violin plots, unfiltered values for percent mitochondrial genes, RNA counts, and RNA features/UMI for the cells of each sample. A red dotted line, filtering cutoff for each respective feature. Bolded samples are those kept after subsequent QC filtering and SAMap comparisons. *, samples removed based on QC cutoffs and DoubleFinder. (C) UMAP, pre-quality control dataset and colored by doublet designation. (D) Sankey plot, correlation of the human and mouse endometrial epithelium. Thickness of connecting lines reflects alignment score.

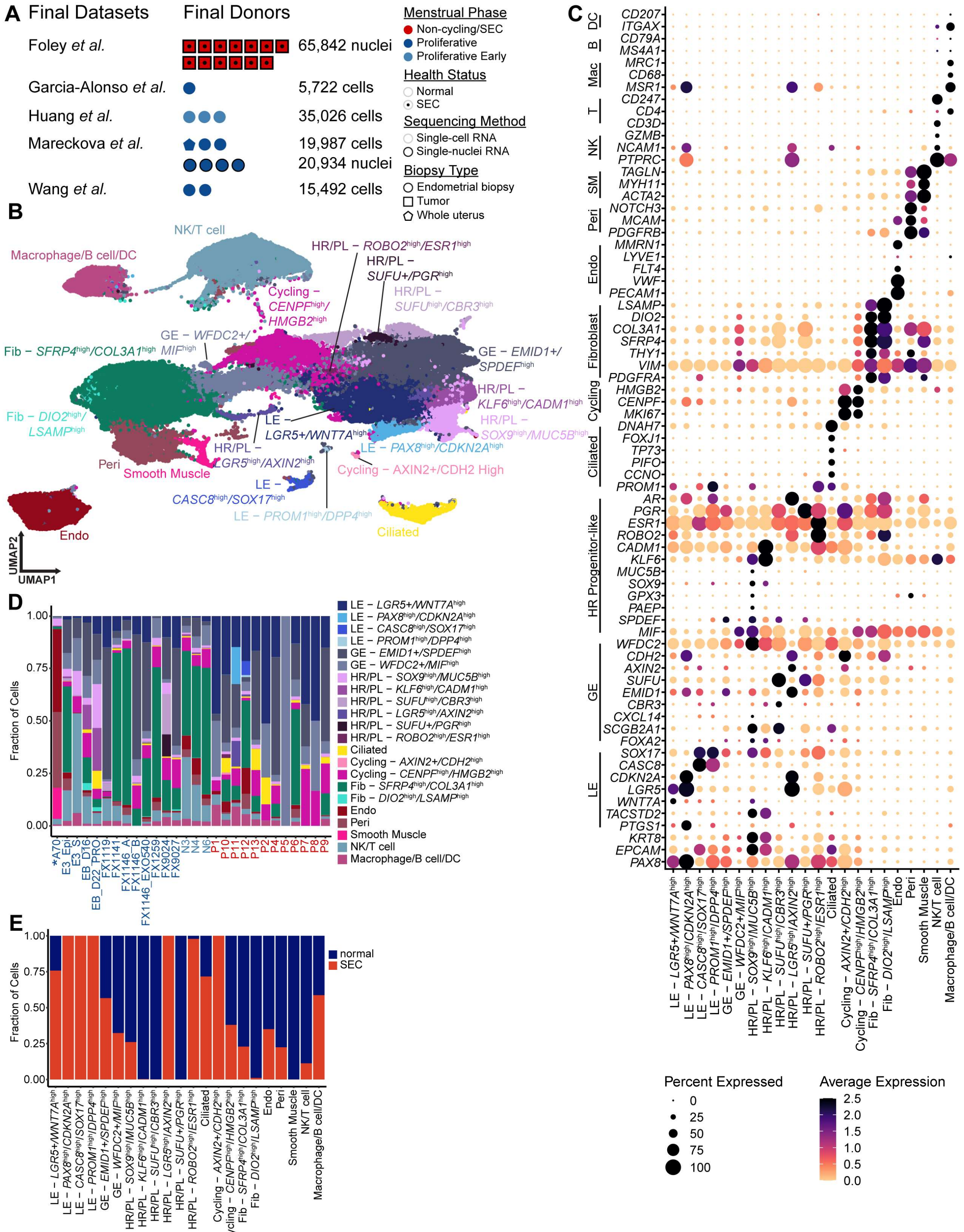

**Supplemental Figure 6. Final human endometrium dataset and cluster naming.** (A) Schematic of the final samples remaining after quality control and sample reduction based on diestrus mouse comparison. (B) UMAP, final human endometrium dataset colored by cluster. (C) Average log-normalized expression of the genes used to name clusters in the final human endometrium dataset. (D) The cell type composition of each human endometrial sample. Sample labels on the x-axis are colored by the patient's menstrual phase. *, whole uterus sample. (E) The contribution of normal versus SEC cells per cluster. LE, Luminal Epithelium; GE, Glandular Epithelium; HR/PL, Hormone-Responsive Progenitor-like; Fib 1-3, Fibroblast; Endo, Endothelium; Peri, Pericyte; DC, dendritic cell.

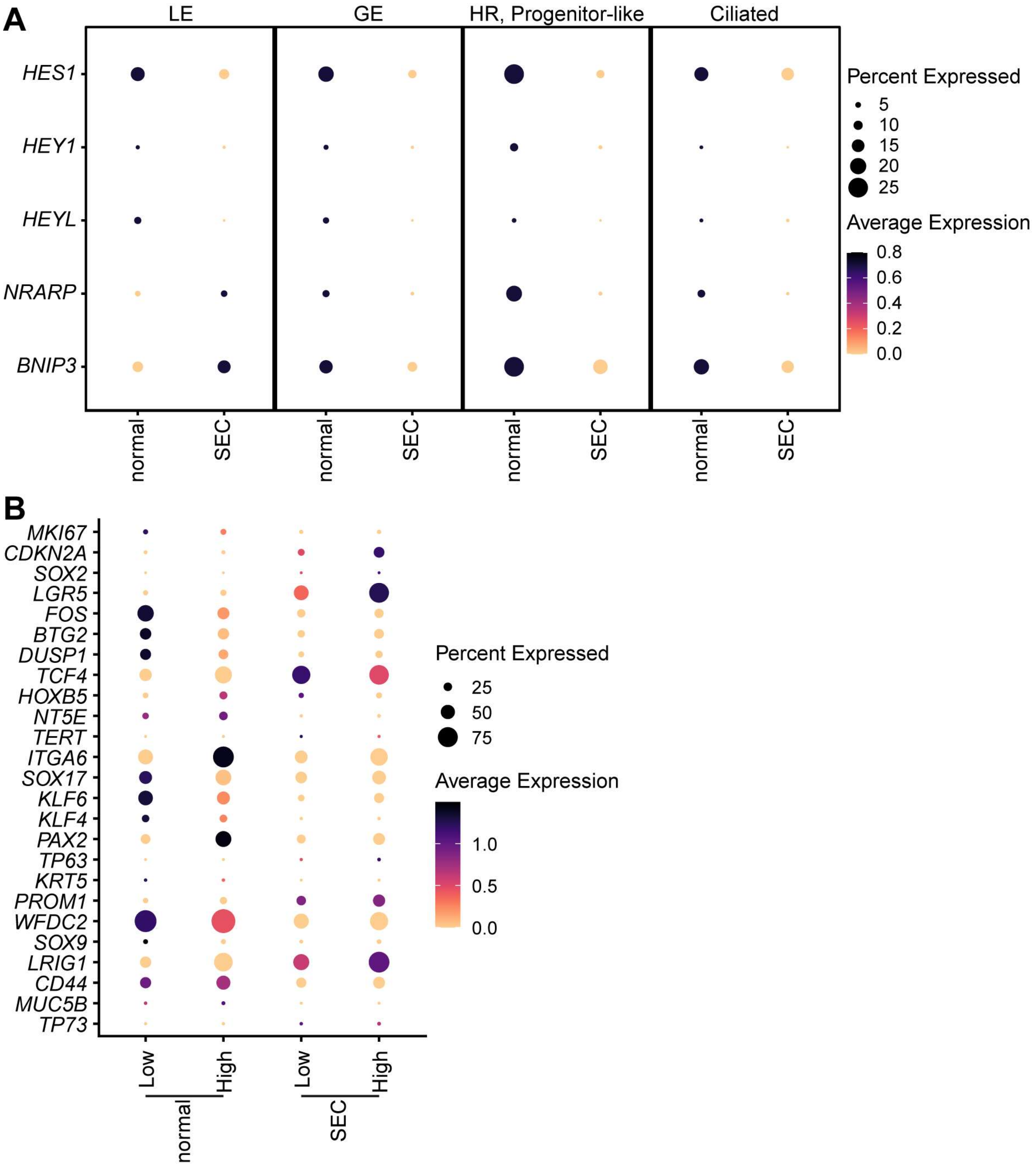

**Supplemental Figure 7. The NOTCH pathway target expression and stem-like changes with human SEC.** (A) Average log-normalized NOTCH target gene expression in the combined LE, GE, Hormone-Responsive (HR) Progenitor-like, and Ciliated populations of human, proliferative uterus samples and human, SEC samples. (B) Average log-normalized stem-like gene expression in the grouped LE, GE, Hormone-Responsive (HR) Progenitor-like, and Ciliated populations in healthy and SEC samples, stratified by *NOTCH2*-expression. Low, below the average normal epithelial expression of *NOTCH2*; High, above the average normal epithelial expression of *NOTCH2*.

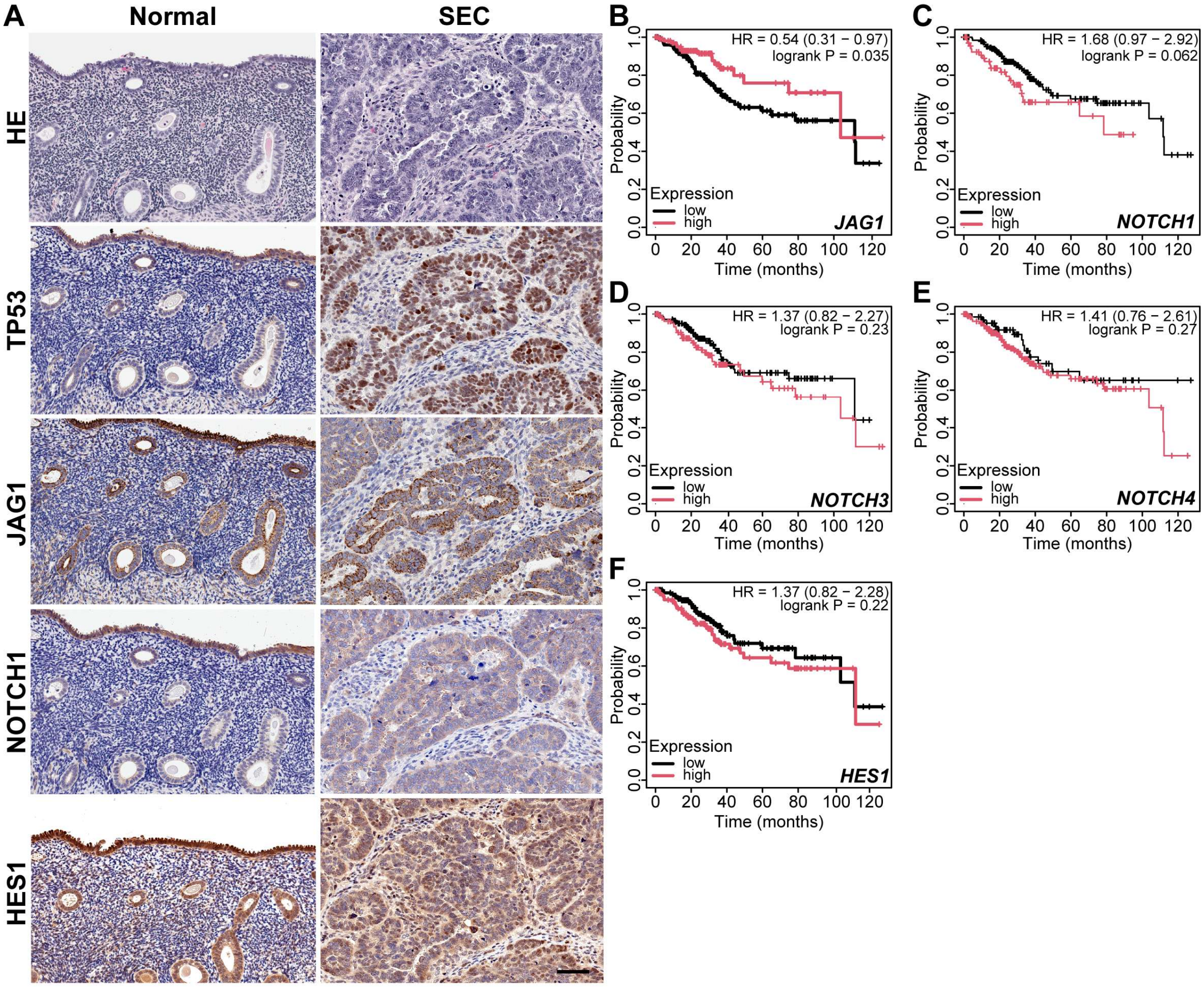

**Supplemental Figure 8. Expression and prognostic associations of NOTCH pathway components in human SEC.** (A) Expression of TP53, JAG1, NOTCH1, and HES1 in the normal human endometrium and serous endometrial carcinoma (SEC). ABC Elite method with hematoxylin counterstaining. Scale bar, 60 µm for all images. (B-F) Kaplan–Meier plots. Overall patient survival stratified by the mRNA expression of *JAG1* (B), *NOTCH1* (C), *NOTCH3* (D), *NOTCH4* (E), and *HES1* (F). Only patients with above median expression of *CDKN2A* were included (*n* = 271).

**Supplemental Tables**

**Supplemental Table 1. Antibodies used for immunostaining.**

| Antigen, conjugation | Antibody source, catalogue number | Clone | Host | Retrieval, Dilution |
|---|---|---|---|---|
| HES1 | Cell Signaling, 11988 | D6P2U | Rabbit | Citrate (30 min) 1:300 (IP-ABC#) |
| JAG1 | Santa Cruz, sc390177 | E-12 | Mouse | Citrate (30 min) 1:1000 |
| Ki67 | Invitrogen, Waltham, MA, USA, 14-5698-82 | SolA15 | Rat | Citrate (10 min), 1:4000 (IP-ABC) |
| NOTCH1 | Cell Signaling, 3608 | D1E11 | Rabbit | Citrate (30 min) 1:300 |
| NOTCH2 | Cell Signaling, 5732S | D76A6 | Rabbit | Citrate (m^, 15 min; h@, 30 min) 1:50 (IF&) 1:100 (IP-ABC) |
| P16 | Sigma, SAB5600077 | RM267 | Rabbit | Citrate (30 min) 1:1600 (IP-ABC) |
| P16 | Abcam, ab241543 | PABLO33B | Rat | Citrate (15 min) 1:50 (IF) |
| P53 | Agilent/Dako, M700101-2 | D07 | Mouse | Citrate (30 min) 1:50 (IF) |
| P53 | Abcam, ab32389 | E26 | Rabbit | Citrate (30 min) 1:400 (IP-ABC) |
| RFP | Rockland, 200-101-379S | PC* | Goat | Citrate (15 min) 1:100 (IF) |
| SOX9 | Abcam, ab185230 | EPR14335 | Rabbit | Citrate (30 min) 1:500 (IP-ABC) |
| Anti-rabbit IgG, biotinylated | Vector Labs, BA-1000-1.5 | PC | Goat | 1:200 |
| Anti-rabbit IgG, Alexa Fluor 594 | Thermo Fisher, A-21207 | PC | Donkey | 1:200 |
| Anti-rat IgG, biotinylated | Vector Labs, BA-4000-1.5 | PC | Rabbit | 1:200 |
| Anti-rat IgG, Alexa Fluor 488 | Vector Labs, A-21208 | PC | Donkey | 1:200 |

*PC, polyclonal; #IP-ABC, immunoperoxidase-ABC Elite method; &IF, immunofluorescence; ^m, mouse tissue; @h, human tissue

**Supplemental Table 2. Single-cell and single-nucleus datasets.**

| | Dataset Name | Description | Analysis used |
|---|---|---|---|
| **Mouse** | Fully Integrated | Includes all mouse cell types and all SEC stages | CellChat; SAMap |
| | Stage-specific All Cells (3 separate datasets) | Epithelial and stromal cells; Normal, pre-dysplastic, and dysplastic samples separated<br>Dryad: https://doi.org/10.5061/dryad.59zw3r2h5 | BayesPrism Reference |
| | Stage-specific Epithelial subset (3 separate datasets) | Epithelial cells only; Normal, pre-dysplastic, and dysplastic samples separated<br>Dryad: https://doi.org/10.5061/dryad.59zw3r2h5 | Label transfer to Full Integrated dataset for CellChat NOTCH pathway analysis; NOTCH component expression; SAMap; BayesPrism reference |
| **Human** | All Menstrual Phase | Includes all human cell types after filtering miscellaneous menstrual phases; normal and SEC samples together; broad cell type labels only | Human menstrual phase correlation to normal, diestrus mouse in SAMap |
| | Final Human | Includes all human cell types; only Proliferative Early, Proliferative, and SEC samples included | Human to mouse cell type correlation; NOTCH pathway component analysis; CellChat |

**Supplemental Table 3. Primers used for PCR.**

| Primer | Type | Use | Sequence |
|---|---|---|---|
| TRE3G-BI_F1 | Forward | N2-ICD cloning | 5' - GGTTAATTAAGGTACCTGGAGTTTACTCCCTATCAGTGATAGAGAACGTATGAAGAG - 3' |
| TRE3G-BI_R1 | Reverse | N2-ICD cloning | 5' - CGTCATGGTCTTTGTAGTCCATGGTGGGCCCCGGTGTCGACTTTACGAGG - 3' |
| mCherry_F1 | Forward | N2-ICD cloning | 5' - CCTAGGCTCGAAATGGTGAGCAAGGGCGAGGAGG - 3' |
| mCherry_R1 | Reverse | N2-ICD cloning | 5' - CCTCCTCGCCCTTGCTCACCATTTCGAGCCTAGG - 3' |
| 3X Flag to EGFP | Forward | N2-ICD cloning | 5' - CGATTACAAGGATGACGATGACAAGATGGTGAGCAAGGGCGAGGAGCTGTTCACC - 3' |
| EGFP to 3X Flag | Reverse | N2-ICD cloning | 5' - GGTGAACAGCTCCTCGCCCTTGCTCACCATCTTGTCATCGTCATCCTTGTAATCG - 3' |
| GAPDH_qPCR_F1 | Forward | RT-qPCR | 5' - CAACTACATGGTCTACATGTT - 3' |
| GAPDH_qPCR_R1 | Reverse | RT-qPCR | 5' - CTCGCTCCTGGAAGATG - 3' |
| N2_ICD_F1 | Forward | RT-qPCR | 5' - CCGAGACTCTAGCAATCACAAG - 3' |
| N2_ICD_R1 | Reverse | RT-qPCR | 5' - GTCCCAGAACCAATCAGGTTAG - 3' |
